# The circadian clock protein REVERBα inhibits pulmonary fibrosis development

**DOI:** 10.1101/781666

**Authors:** Peter S. Cunningham, Peter Meijer, Alicja Nazgiewicz, Simon G. Anderson, Lee A. Borthwick, James Bagnall, Gareth B. Kitchen, Monika Lodyga, Nicola Begley, Rajamayier V. Venkateswaran, Rajesh Shah, Paul F. Mercer, Hannah J. Durrington, Neil C. Henderson, Karen Piper-Hanley, Andrew J. Fisher, Rachel C. Chambers, David A. Bechtold, Julie E. Gibbs, Andrew S. Loudon, Martin K. Rutter, Boris Hinz, David W. Ray, John F. Blaikley

## Abstract

Pulmonary inflammatory responses lie under circadian control; however the importance of circadian mechanisms in fibrosis is not understood. Here, we identify a striking change to these mechanisms resulting in a gain of amplitude and lack of synchrony within pulmonary fibrotic tissue. These changes result from an infiltration of mesenchymal cells, an important cell type in the pathogenesis of pulmonary fibrosis. Mutation of the core clock protein REVERBα in these cells exacerbated the development of bleomycin-induced fibrosis, whereas mutation of REVERBα in club or myeloid cells had no effect on the bleomycin phenotype. Knockdown of REVERBα revealed regulation of the poorly described transcription factor TBPL1. Both REVERBα and TBPL1 altered integrinβ1 focal adhesion formation, resulting in increased myofibroblast activation. The translational importance of our findings was established through analysis of two human cohorts. In the UK Biobank circadian strain markers (sleep length, chronotype and shift work) are associated with pulmonary fibrosis making them novel risk factors. In a separate cohort REVERBα expression was increased in human idiopathic pulmonary fibrosis (IPF) lung tissue. Pharmacological targeting of REVERBα inhibited myofibroblast activation in IPF fibroblasts and collagen secretion in organotypic cultures from IPF patients, suggesting targeting REVERBα could be a viable therapeutic approach.

**Significance:** The circadian clock plays an essential role in energy metabolism, and inflammation. In contrast the importance of the clock in the pathogenesis of fibrosis remains poorly explored. This study describes a striking alteration in circadian biology during pulmonary fibrosis where the relatively arrhythmic alveolar structures gain circadian but desynchronous rhythmicity due to infiltration by fibroblasts. Disruption of the clock in these cells, which are not widely implicated in circadian pathophysiology, results in a pro-fibrotic phenotype. Translation of these findings in humans revealed previously unrecognised important circadian risk factors for pulmonary fibrosis (sleep length, chronotype and shift work). In addition, targeting REVERBα repressed collagen secretion from human fibrotic lung tissue making this protein a promising therapeutic target.

## Introduction

The lung is highly circadian, resulting in temporal gating of a number of inflammatory (1–3) and anti-oxidant responses (4). Local timing is dominated by non-ciliated, bronchial epithelial cells (club cells) (2) and alveolar macrophages (3, 5). In contrast, alveolar structures typically show only weak circadian oscillations (6), with implications for the pathogenesis of diseases such as pulmonary fibrosis. Despite this, genetic disruption of the *Clock* gene (4), impairing circadian oscillations, exaggerates early mouse pulmonary responses to bleomycin challenge; a model of pulmonary fibrosis (7).

Pulmonary fibrosis, including idiopathic pulmonary fibrosis (IPF), is frequently fatal with existing treatments slowing progression rather than curing the disease (8). The causes and non-genetic risk factors for IPF are poorly understood, with several studies implicating age, sex, smoking and more recently air pollution (9). IPF is characterized histologically by the development of fibroblastic foci in the lung parenchyma (10). Cells in these foci are typically activated myofibroblasts (11) derived from multiple sources (12, 13), including pulmonary fibroblasts and pericytes (11, 14). Myofibroblasts secrete collagen resulting in abnormal lung function and are characterized by increased focal adhesion formation and acquisition of a contractile cytoskeleton with alpha smooth muscle actin (αSMA)-positive stress fibers (15). In addition to fibroblasts, pulmonary fibrosis involves other cell types e.g. club cells (9) and macrophages (16) regulating the accumulation of fibroblasts and therefore the deposition of the extracellular matrix. As these cell types maintain autonomous circadian oscillations (2, 5), examination of circadian factors and mechanisms in the pulmonary fibrotic response is warranted.

The circadian clock is an internal timing mechanism (17), allowing temporal segregation of both pathophysiological and physiological programs (18, 19). At the cellular level the circadian clock consists of a transcription-translation feedback loop (20), in which the positive elements CLOCK and BMAL1 drive expression of two negative feedback arms controlled by PERIOD/CRYPTOCHROME (PER/CRY) and the two paralogues REVERBα and REVERBβ. In turn these negative feedback arms repress BMAL1/CLOCK heterodimer transactivation function (PER/CRY), or BMAL1 expression (REVERBα/β). The resulting 24-hour oscillations in protein expression can be disrupted through environmental disruption (e.g. shift-work schedules), or genetic deletion of core clock components producing inflammatory and metabolic phenotypes (5, 21, 22).

Here, we show that fibrotic mouse lungs exhibited amplified, but desynchronous circadian rhythms, with a dominant role for myofibroblasts. Disruption of the core clock protein REVERBα in fibroblastic cells resulted in exaggerated pulmonary fibrotic response to bleomycin in mice. In culture, REVERBα knockdown resulted in increased myofibroblast differentiation via the transcription factor TBPL1, through altering the formation of integrinβ1 focal adhesion expression. Furthermore exposure to circadian stresses such as late chronotype, shift work, and altered sleep duration are all associated with idiopathic pulmonary fibrosis and clock gene expression is altered in IPF versus normal human lung. Targeting of REVERBα by a synthetic ligand repressed myofibroblast differentiation and collagen secretion in cultured fibroblasts and lung slices obtained from patients with lung fibrosis.

## Results

### Myofibroblasts drive high amplitude, but desynchronous circadian oscillations in fibrotic lung

Precision cut lung slices from transgenic mPER2∷LUC mice (2) were used to track circadian oscillations in real-time after bleomycin-induction of fibrosis (Fig. 1A, S1A,B, Video S1). Fibrotic areas were identified by loss of lung architecture in the brightfield image, and confirmed with increased collagen deposition when the slices were fixed for histology (Fig. S1A,B). The amplitude of PER2 oscillations in the fibrotic areas was increased compared to non-fibrotic parenchyma lung (Fig. 1A-B,D). The fibrotic parenchyma also had greater phase desynchrony compared to regions in the non-fibrotic parenchyma (Fig. 1C). One possible explanation for these changes is the cell density in the fibrotic parenchyma. To explore this precision cut lung sections (PCLS) were stained with Hoechst. There was a greater intensity of staining in fibrotic areas compared to non-fibrotic areas, but this did not correlate with bioluminescence (Fig. S1C-E). Another possible explanation is infiltration by a more rhythmic cell type, therefore we deleted the essential core clock component BMAL1 (23) in both fibroblasts and club cells to ablate cell-autonomous rhythms. BMAL1 deletion in club cells (CCSP-Bmal1), the main oscillatory cells in the lung (6), had no effect on the increased amplitude seen in fibrotic regions (Fig. 1D,G, S1G, Video S2). In contrast, BMAL1 deletion in pericyte/fibroblast lineage (Pdgfrb-Bmal1^−/−^) restored the amplitude of lung oscillations in fibrotic lung to levels measured in unaffected lung tissue (Fig. 1E-G, S1F, Video S3).

**Figure 1.**
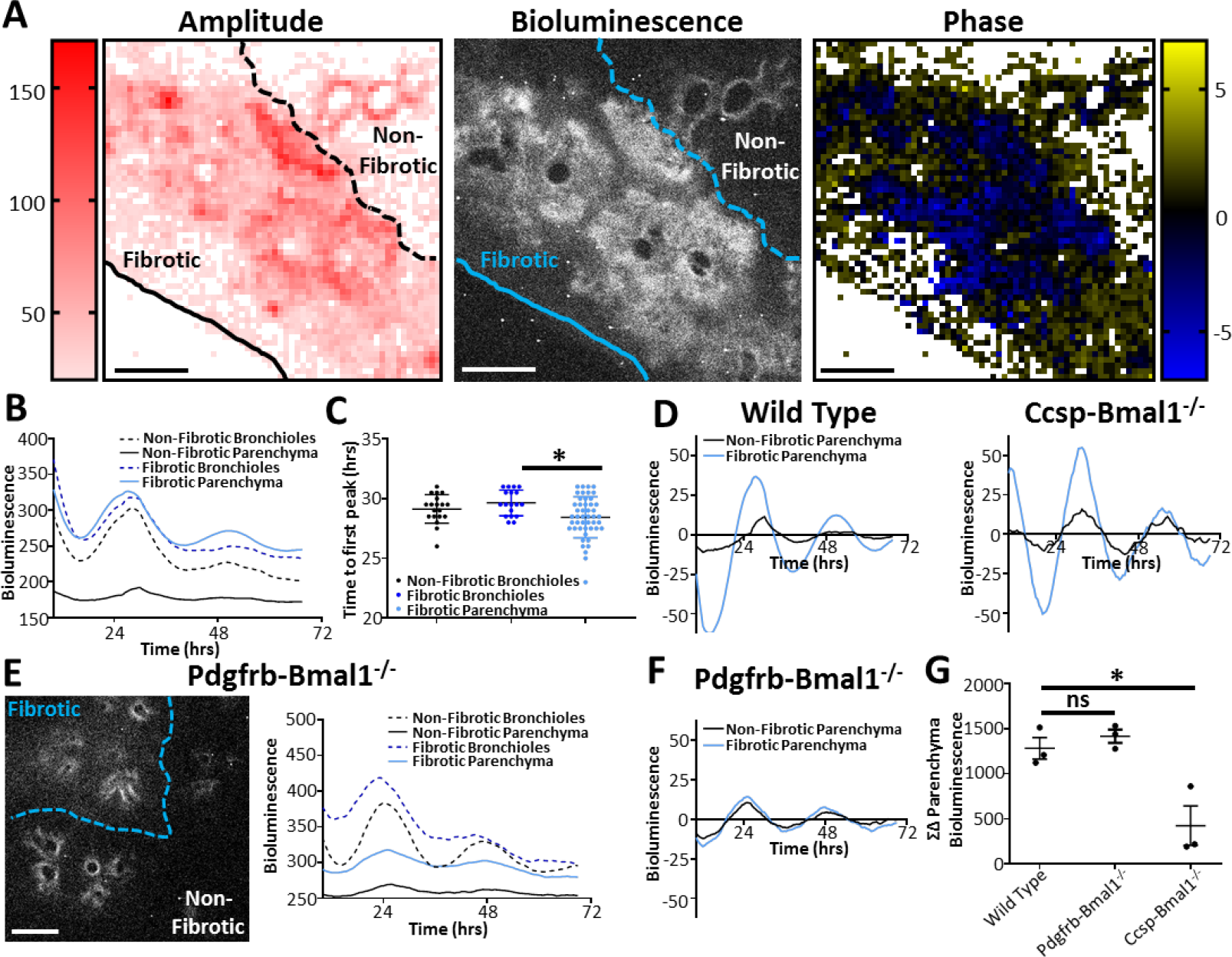
Desynchronous circadian oscillations occur in Pulmonary Fibrosis. **A)** Bioluminescent image, along with heatmaps of amplitude and phase taken from the same precision cut lung slice obtained from a mPER2∷luc mouse fourteen days after *in vivo* bleomycin (3U/kg); scale bar, 500μm (data is representative of three separate experiments) **B)** Bioluminescent intensity plotted against time for both parenchyma and bronchioles in fibrotic and non-fibrotic regions (data is representative of three separate experiments) **C)** Time to first peak for bronchioles and parenchyma in fibrotic and non-fibrotic areas (**=p<0.01, ANOVA with post-hoc Dunnett’s test using 18, 19 and 48 representative sections for healthy airways, fibrotic airways and fibrotic parenchyma respectively in the lung slice, data is representative of three separate experiments) **D)** Bioluminescent intensity plotted against time (24hr moving average baseline subtracted) for the representative slices shown in (A) and Ccsp-Bmal1^−/−^ mice shown in Fig. S1G **E)** Representative bioluminescent image along with bioluminescent intensity plotted against time for a precision cut lung slice fourteen days after *in vivo* bleomycin treatment (image and data representative of 3 mice in 3 separate experiments) in the Pdgfrb-Bmal1^−/−^ mPER2∷luc mouse; scale bar, 500μm **F)** Bioluminescent intensity plotted against time (24hr moving average baseline subtracted) for the Pdgfrb-Bmal1^−/−^ representative slice shown in (E) **G)** Difference in bioluminescence between fibrotic and non-fibrotic parenchyma over 3 days in precision cut lung slices from WT, Ccsp-Bmal1^−/−^ and Pdgfrb-Bmal1^−/−^ mice after *in vivo* bleomycin treatment (*=p<0.05, ANOVA post-hoc Dunnett’s n=3 mice done in three separate experiments per condition).

To test if pro-fibrotic factors are capable of modifying circadian signals between cells, lung slices and fibroblasts were treated with TGFβ, and inflated to mimic changes in the cellular mechanoenvironment (Fig. S2A). TGFβ induced changes in circadian phase, with the magnitude of effect being dependent on concentration and phase (Fig. S2B,C). Lung inflation also increased the amplitude of the PER2∷LUC oscillation (Fig. S2D).

### REVERBα in fibroblasts suppresses the development of pulmonary fibrosis

REVERBα is an orphan nuclear receptor, and both an essential core clock factor, and major clock output pathway. Its function can be disrupted by deletion of its DNA binding domain, and small molecular ligands are available to modulate activity. Therefore we deleted the REVERBα DNA binding domain (Fig. 2A), under Pdgfrb control (24). This resulted in an exaggerated fibrotic response (Fig. 2B,C) and increased accumulation of αSMA-positive myofibroblasts in response to bleomycin (Fig. 2D,E). Wild-type and transgenic mice did not differ in lung parameters following saline inoculation (Fig. 2B,E, S3A-B). Importantly, REVERBα genetic disruption in myelomonocytic cells, or bronchial epithelial cells did not affect the development of the fibrotic phenotype (Fig. S3C,D).

**Figure 2.**
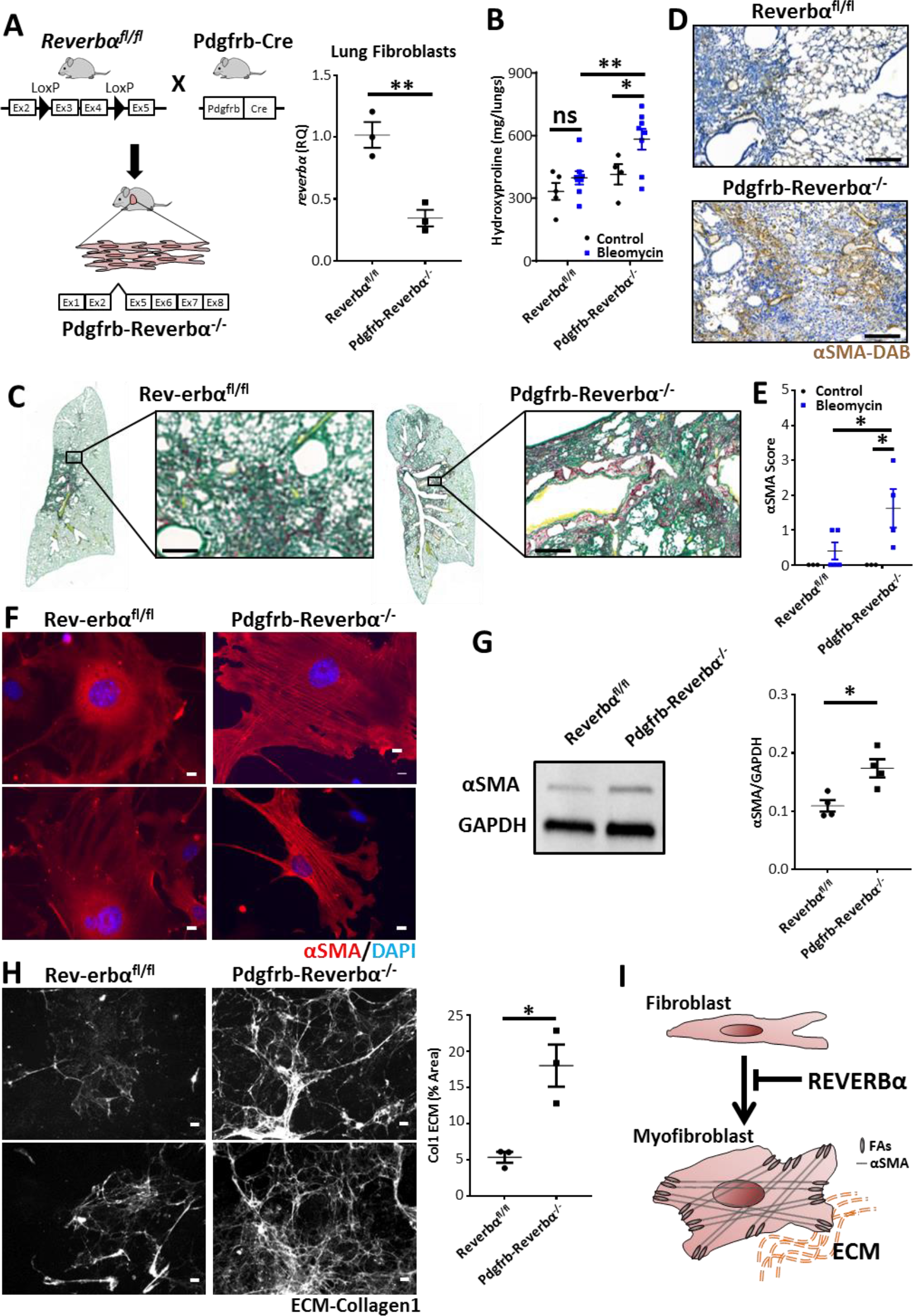
REVERBα alters susceptibility to pulmonary fibrosis through its effect on myofibroblast differentiation. **A)** Schematic showing generation of Pdgfrb-Reverbα^−/−^ mice combined with qPCR analysis of *reverbα* expression in lung fibroblasts (n=3 animals, **=p<0.01, Student’s t-test) **B)** Hydroxyproline measurement in lungs from Pdgfrb-Reverbα^−/−^ mice and littermate controls 28 days following challenge with intra-tracheal bleomycin (2U/kg) or saline (n=4-5 saline and 8 bleomycin per genotype,*=p<0.05, **=p<0.01, 2-way ANOVA Holm-Sidak post hoc test) **C)** In a separate experiment histology (Picrosirius red) of lungs was examined 28 days following challenge with intra-tracheal bleomycin (representative image from 4 animals treated with bleomycin per genotype scale bar=200μm) **D)** Immunohistochemical staining of myofibroblasts (anti-αSMA, 3,3′-Diaminobenzidine (DAB)) from Pdgfrb-Reverbα^−/−^ mice and littermate controls 28 days following intra-tracheal bleomycin challenge (representative image from 4 animals treated with bleomycin per genotype scale bar=200μm) **E)** Histological scoring (Grade 0-4) for the presence of αSMA staining 28 days following intra-tracheal bleomycin challenge (n=3 saline and 4-5 bleomycin per genotype,*=p<0.05, 2-way ANOVA Holm-Sidak post hoc test **F)** Representative immunofluorescence images of primary lung fibroblast cultures from naïve Pdgfrb-Reverbα^−/−^ mice and littermate controls showing intra-cellular αSMA (red) (n=3 animals per genotype; scale bar=10μm) combined with a **G)** representative immunoblot and quantification of intracellular αSMA from primary lung fibroblast cultures (n=4 animals per genotype, **=p<0.01, Student’s t-test) **H)** Representative collagen-1 ECM images and quantification following culture of Pdgfrb-Reverbα^−/−^ and Reverbα^fl/fl^ primary lung fibroblasts (n=3 animals per genotype, **=p<0.01, Student’s t-test; scale bar, 50μm) **I)** Schematic illustrating the action of REVERBα in inhibiting fibroblast/myofibroblast differentiation.

Characterization of primary fibroblasts explanted from Pdgfrb-Reverbα^−/−^ lungs *ex vivo* revealed increased expression of αSMA and increased secretion of collagen-1, markers of myofibroblast activation (Fig. 2F-H). This indicates a fibroblast-intrinsic change driven by disruption of REVERBα, with culture on hard plastic providing the environmental trigger for initiation of the myofibroblast differentiation program (Fig. 2I).

### Knockdown of REVERBα *in vitro* enhances myofibroblast activation through the transcription factor TBPL1

Next, we set out to identify REVERBα gene targets using siRNA knockdown of REVERBα in both mouse and human lung fibroblast cell lines (Fig. S4A). REVERBα knockdown resulted in myofibroblast activation in lung fibroblast cells (Fig 3A-B, S4B-D). Although many genes were regulated by REVERBα knockdown, only two were repressed at both timepoints (12&24 hours) and in both cell lines (Fig. 3C, S4E,F). One was *PLOD2*, a proline hydroxylase required for collagen processing. The second was *TBPL1*, a poorly characterized transcription factor. As the role of PLOD2 in collagen processing is already well characterized, we turned to *TBPL1*, and verified loss of protein expression with REVERBα knockdown (Fig. 3D). Knockdown of TBPL1 caused a similar induction of αSMA expression to that seen with REVERBα knockdown (Fig. 3E) suggesting that REVERBα and TBPL1 contribute to the same pathway.

**Figure 3.**
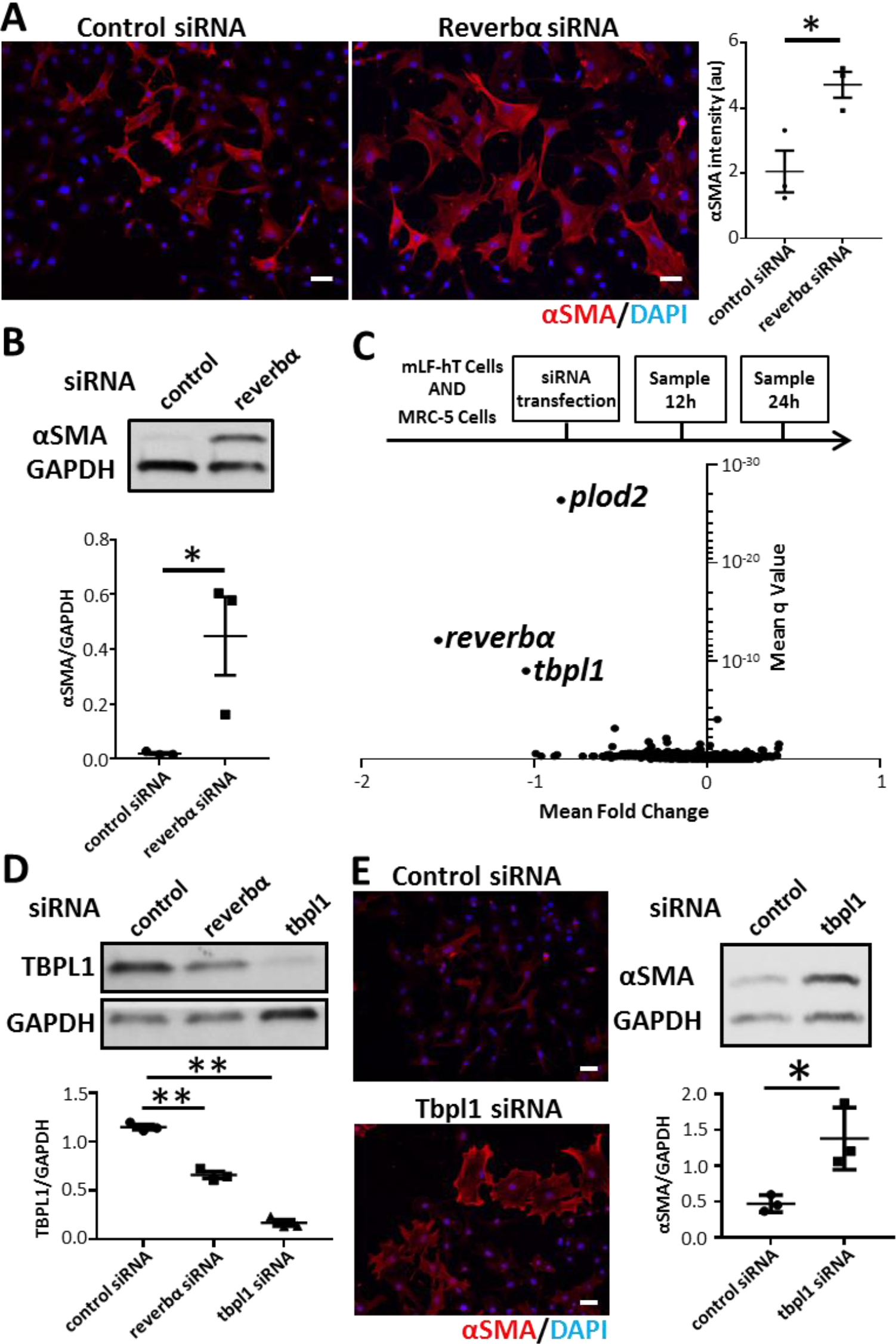
REVERBα alters myofibroblast differentiation via TBPL1. **A)** Immunofluorescent staining and quantification for the myofibroblast marker αSMA after control (non-targeting) or *Nr1d1* (REVERBα) siRNA knockdown in mLF-hT cells, scale bar, 50μm (*=p<0.05, student’s t-test, n=3 separate transfections) **B)** Immunoblot and densitometry for αSMA in MRC-5 cells after control (non-targeting) or *REVERBα* siRNA knockdown (representative immunoblot shown, n=3 separate transfections, *=p<0.05 Student’s t-test) **C)** Schematic of RNA-seq sample preparation. Control (non-targeting) or *Reverbα* siRNA knockdown was performed in two fibroblast cell lines (mLF-hT cells and MRC-5). Samples were collected for RNA-seq analysis 12 and 24 hours after siRNA transfection from three separate transfections per timepoint and cell line. Pooled analysis of all four different RNA-seq experimental conditions shown by a volcano plot (mean fold change plotted against mean q-value) **D)** Immunoblot of TBPL1 following control (non-targeting), *Reverbα* or *Tbpl1* siRNA knockdown in mLF-hT cells (representative immunoblot shown, n=3 separate transfections, **=p<0.01 Student’s t-test) **E)** Immunofluorescence and immunoblotting for αSMA after control (non-targeting) or *Tbpl1* siRNA knockdown in mLF-hT cells (representative immunoblot shown, n=3 separate transfections *=p<0.05 Student’s t-test); scale bar, 50μm

### REVERBα and TBPL1 regulate Integrinβ1 expression

To decipher how REVERBα and/or TBPL1 suppress myofibroblast activation in fibrotic lungs and in the stiff cell culture environment, we focused on focal adhesions, crucial mechanotransducing elements that control myofibroblast activation (25). Knockdown of either REVERBα or TBPL1 resulted in increased sizes and numbers of vinculin/tensin1 positive focal adhesion complexes (Fig. 4A, S5A-C). This increase in size suggests progression to the super-mature focal adhesions involved in myofibroblast differentiation (25). In contrast, overexpression of REVERBα or TBPL1 caused the opposite effect (Fig. 4B, S5D,E). Integrinβ1, the common subunit of all collagen1-binding integrins, has previously been linked to myofibroblast activation in the liver (26), lung (27) and scleroderma (28). Knockdown of either REVERBα or TBPL1 resulted in an increase in both size and number of integrinβ1-positive focal adhesion complexes (Fig. 4A, S5A). Furthermore, knockdown of integrinβ1 prevented the induction of αSMA seen in fibroblasts cultures subjected to REVERBα knockdown (Fig. 4C,D), highlighting the requirement for integrinβ1 for REVERBα-mediated myofibroblast activation (Fig. 4E).

**Figure 4.**
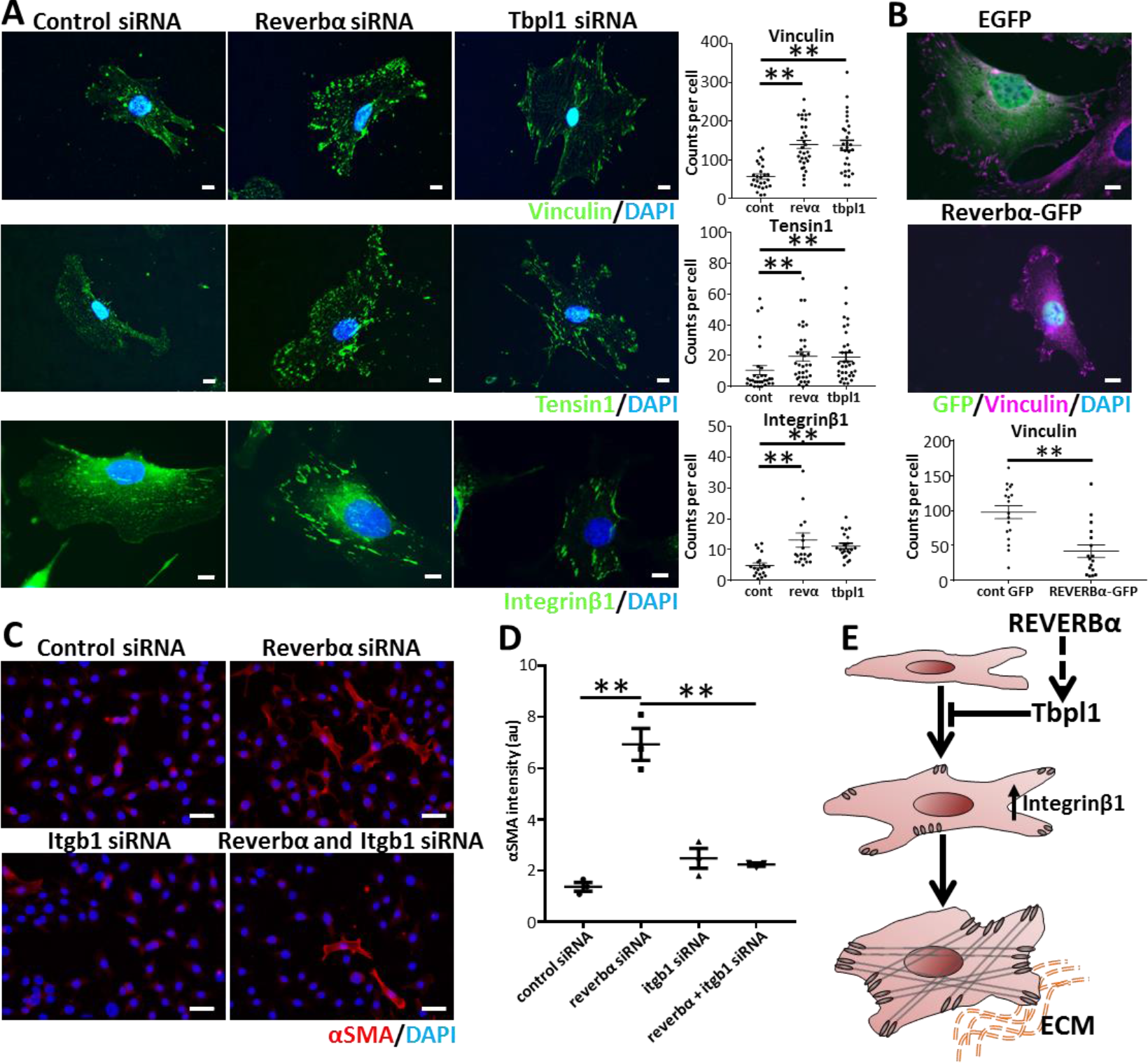
REVERBα and TBPL1 affect myofibroblast differentiation through changes in integrinβ1 expression. **A)** Representative immunofluorescent images and quantification per cell of vinculin, tensin1 and integrinβ1 following siRNA knockdown of *Reverbα* or *Tbpl1* compared to control (non-targeting) siRNA in mLF-hT cells (n=3 transfections, **= p<0.01 ANOVA post-hoc Dunnett’s •= individual cells from 3 transfections; scale bar, 10μm, cont = control siRNA, revα = *Reverbα* siRNA **B)** Representative immunofluorescence image after mLF-hT cells have been transfected with REVERBα-GFP plasmid or an empty-GFP plasmid. Cells were stained for GFP, Vinculin and nuclei (DAPI) (n=3 separate transfections; scale bar, 10μm with the focal adhesion number being quantified per cell (n=3 transfections, **= p<0.01, Student’s t-test •= individual cells from 3 transfections **C)** Representative immunofluorescence images; scale bar, 50μm and **D)** Quantification of the myofibroblast marker αSMA, using immunofluorescence, following dual siRNA knockdown (control or *Reverbα* in the presence or absence of *Itgb1*) in mLF-hT cells (n=3 separate transfections, *p=<0.05 ANOVA post-hoc Dunnett’s) along **E)** Schematic demonstrating how both REVERBα and TBPL1 regulate integrinβ1 which in turn affects myofibroblast differentiation.

### Circadian factors are associated with pulmonary fibrosis in humans

Several human factors have been associated with circadian, or sleep-deprivation strain, including evening chronotype, shift work and sleep duration. We therefore investigated whether these factors were associated with pulmonary fibrosis in the UK Biobank (29) (n=500,074). Following adjustment for known risk factors for pulmonary fibrosis (BMI, smoking, sex and age) short or long sleep duration (<7h or >7h) were associated with pulmonary fibrosis (Fig. 5A), with the size of the odds ratio being greater than the established risk factors of age, sex or smoking in the multivariable model. Shift work (OR 1.353 95%CI 1.069-1.710) and evening chronotype (OR 1.040 95%CI 1.001-1.080) were also associated with pulmonary fibrosis (Tables S1-6) by a smaller degree, however this is comparable to other diseases were these variables are risk factors (30–32).

**Figure 5.**
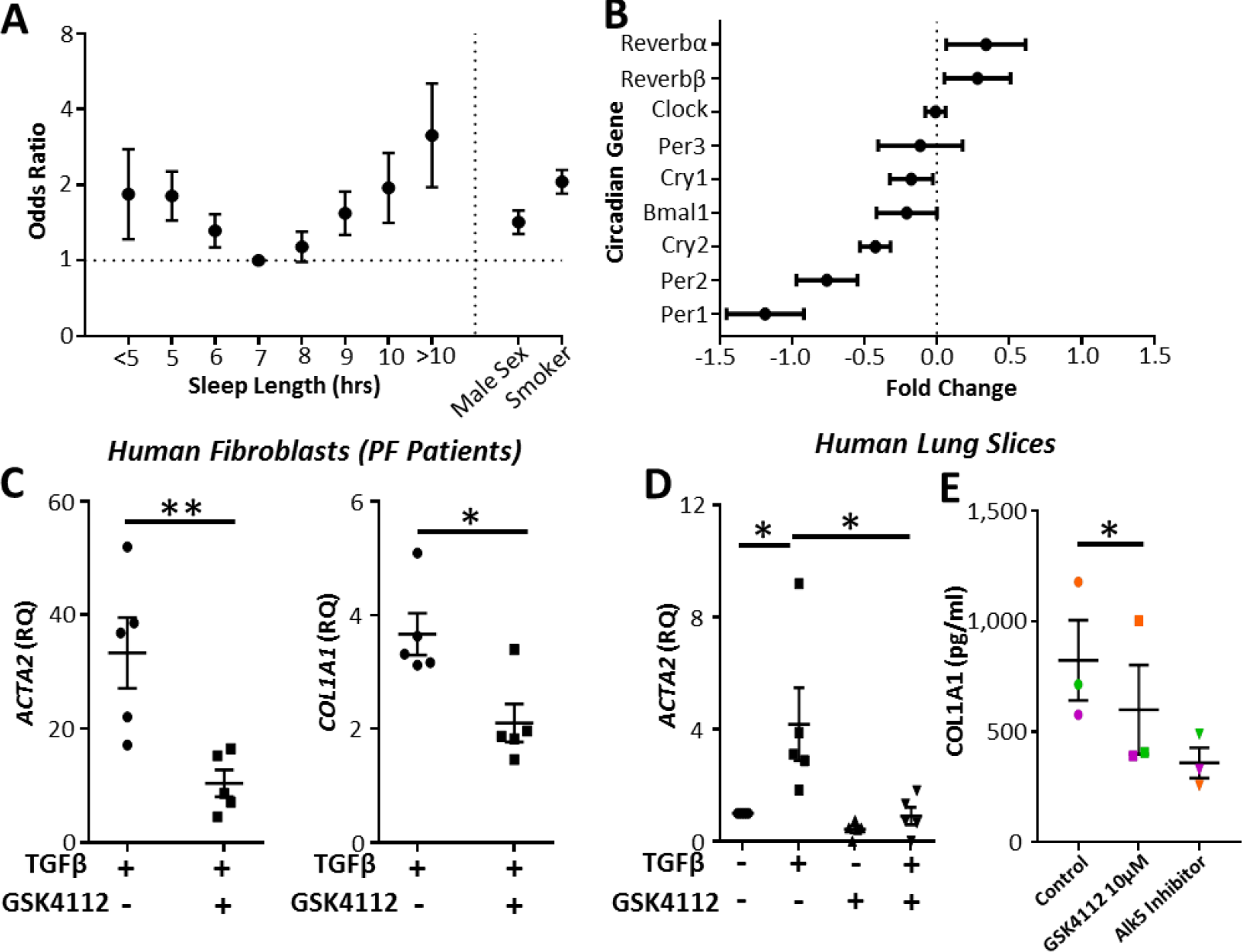
Circadian factors are associated with idiopathic pulmonary fibrosis, where a REVERB ligand represses collagen secretion. **A)** Odds ratios for the association between pulmonary fibrosis and sleep duration (OR ± 95% confidence interval, logistic regression n=500,074 subjects from the UK biobank) **B)** Changes in clock gene expression in idiopathic pulmonary fibrosis compared to control subjects from a previously published genome array (GSE 47460) (fold change ± 95% confidence interval, n=90 controls and 98 patients with IPF) **C)** qPCR for αSMA (*acta2*) and Collagen1 (*col1a1*) following TGFβ-stimulation (2ng/ml) in primary human lung fibroblasts obtained from patients with pulmonary fibrosis in the presence or absence of GSK4112 (10μM). (n=5 fibrotic patients, *=p<0.05, **=p<0.01 Student’s t-test,) **D)** qPCR for αSMA (*acta2*) expression following treatment with TGFβ (2ng/ml) and GSK4112 (10μM) in human precision cut lung slices (n=5 patients, *=p<0.05 Student’s t-test) **E)** ELISA analysis of secreted collagen-1 in TGFβ stimulated precision cut lung slices obtained from three patients with idiopathic pulmonary fibrosis treated with the REVERB ligand GSK4112 (10μM) and an Alk5 inhibitor (1μM) as positive control (n=3 *=p<0.05, Paired Student’s t-test).

### Disordered clock gene expression occurs in idiopathic pulmonary fibrosis

To look for evidence of circadian clock disruption in idiopathic pulmonary fibrosis (IPF), we analysed lung gene expression in a previously published microarray from the lung research consortium (33). Comparison with normal lung revealed significant differences in *PER1/2*, *CRY 2* and *REVERBα/β* (Fig. 5B), all encoding components of the negative feedback arm of the core circadian clock. In addition, *TBPL1* was upregulated in pulmonary fibrosis, correlating with *REVERBα* expression (Fig. S4G).

### A REVERB ligand inhibits myofibroblast differentiation and represses collagen secretion in tissue from pulmonary fibrotic patients

Finally, we tested whether a REVERBα ligand could repress pulmonary fibrosis. The well-characterized REVERBα agonist GSK4112 (34) repressed TGFβ-induced expression of αSMA (*ACTA2*) and collagen-1 (*COL1A1*) in primary human lung fibroblasts from patients with pulmonary fibrosis (Fig. 5C). Similarly, TGFβ induction of αSMA and collagen 1 transcription was prevented by GSK4112 treatment in precision-cut human lung, organotypic slice cultures from healthy control subjects (Fig. 5D). Finally we studied the effects of this ligand in PCLS from IPF patients along with an Alk5 inhibitor; known to inhibit Col1a1 secretion (35). GSK4112 repressed Col1a1 secretion (Fig. 5E) in a similar manner to the Alk5 inhibitor.

## Discussion

Pulmonary fibrosis is an intractable and fatal disease. We have previously identified the lung as a highly circadian organ, and that responses to environmental insults are regulated and shaped by the circadian clock. Therefore we analysed mouse lung fibrosis, finding newly-emergent, and strong circadian oscillations driven by fibroblasts. The prevalent pro-fibrotic growth factor TGFβ was capable of transmitting timing information to recipient cells, and disruption to the core circadian clock in fibroblasts increased fibrotic response to bleomycin instillation. *In vitro* analysis identified a circuit linking the core clock through REVERBα, to TBPL1, and the focal adhesions important for myofibroblast activation. In human IPF lung tissue pharmacological targeting of the clock impacted a surrogate measure of fibrotic progression, and we found a new association between sleep duration, which is a product of the circadian clock, and risk of pulmonary fibrosis.

Several studies have found that circadian responses in the lung are gated through club cells (2) or macrophages (5). A previous report suggested that the acute inflammatory phase (7 days) of the bleomycin response lay under circadian control (4), therefore, we investigated circadian function in developing fibrosis. Surprisingly, there were higher amplitude circadian oscillations in fibrotic tissue compared to normal lung tissue but these oscillations were desynchronous, suggesting a possible role for circadian mechanisms (36). The process of fibrosis involves several different cell types including club cells, macrophages and fibroblasts (9), but as genetic deletion of the only non-redundant circadian gene *Bmal1* to the pericyte lineage stopped the emergent oscillations the importance of fibroblasts was established. The importance of the fibroblast was further confirmed by finding that REVERBα deletion in these cells impacted the fibrotic response, but disruption in other cell types was without effect. It is already known that circadian oscillations in fibroblasts are robust (37), altering wound-healing (38, 39).

Mechanistically, knockout or knockdown of REVERBα promoted myofibroblast activation *in vitro*, with the reverse effects seen with REVERBα overexpression. Analysis of REVERBα gene targets revealed striking enrichment for a single transcription factor, TBPL1, and the emergence of a coherent pathway converging on increased formation of integrinβ1 focal adhesion complexes. To the best of our knowledge TBPL1 has not been previously implicated in fibrotic disease, but we found its expression elevated in human IPF tissue. This and the elevated REVERBα expression is an apparent paradox, as both proteins inhibit myofibroblast activation. Therefore, we hypothesize that the increase in both TBPL1 and REVERBα in fibrotic tissue results from tissue compensation in response to fibrosis, making it a promising therapeutic pathway. Integrinβ1, emerged as the final effector, and the focal adhesions associated with it have previously been established (26) to be important for myofibroblast activation.

We have successfully used large-scale human cohorts, such as the UK Biobank, to explore connections between measures of circadian strain (shiftwork, chronotype, and sleep) and prevalent disease (32, 40, 41). Low prevalence diseases such as pulmonary fibrosis present unique challenges. To address this we identified people with pulmonary fibrosis participating in the UK Biobank (29), and linked them with information from Hospital Episode Statistic data (42). Importantly, patients were not screened for pulmonary fibrosis on enrolling in the Biobank therefore we cannot comment on causality, but it is clear that short sleep length is associated with pulmonary fibrosis and this is as least as strong as existing risk factors for this disease (43) indicating potential clinical relevance. An association with long sleep duration was also found which may be biological (44) or due to confounders (45).

We, and others, have developed tool compounds capable of activating REVERBα (46, 47). These permit extension of our studies to primary human tissue, which is hard to genetically manipulate. Here, we show a marked inhibition of the myofibroblast phenotype, blunted fibrotic response to TGFβ stimulation and reduced collagen-1 secretion in IPF precision cut lung slices. We, and others, have shown that these compounds have off target effects (46, 48), therefore it is reassuring that knockdown and overexpression of REVERBα in human fibroblasts had similar effects to both our mice studies and also the ligand. The recent publication (48) that the only ligand with suitable PK for *in vivo* experiments has significant off target effects combined with the lack of translation from the mouse bleomycin model to the clinic (49) precludes an *in vivo* mouse experiment to confirm its therapeutic effectiveness. Furthermore tissue from IPF patients is emerging as a predictive model for human translation.

Taken together our results identify a surprising, and potent role for the core circadian clock factor REVERBα in the activation of myofibroblasts via a novel pathway incorporating a poorly characterised transcription factor TBPL1 which affects the development of pulmonary fibrosis.

## Methods

### Mouse Lines

mPER2∷luc transgenic mice were previously described (50). The Rev-erbα^fl/fl^ mouse (Rev-erbαDBD^m^) and Cre drivers targeting club cells (CCSP^icre^) and myeloid cells (Lysm^cre^) are as previously described (1). The PDGFRβ^cre^ mouse was a kind gift from Henderson and has been previously described (14). The Bmal1^fl/fl^ mouse has been previously described (2).

### Cell Culture

MRC-5 cells or mLF-hT cells (16) were cultured in DMEM media.

### *In vivo* Bleomycin

Male mice were challenged intratracheally with Bleomycin (sigma) or Saline (vehicle).

### Bioluminescence Microscopy

Precision-cut organotypic lung slices (PCLS) were prepared as described before (2). Bioluminescence images were obtained using a 2.5× objective (Zeiss) and captured using a cooled Andor iXon Ultra camera over a 30 minute integration period.

### Immunofluorescence

For αSMA staining, cells in 35mm dishes were fixed in 4% PFA/0.2% Triton X, followed by ice cold methanol fixation. For focal adhesions proteins cells were exposed to ice-cold cytoskeleton buffer (51) for 10 minutes followed by 4% PFA fixation for a further 10 minutes.

### RNA-seq

siRNA transfected mLF-hT and Mrc5 cells were lysed and RNA was extracted using the ReliaPrep RNA miniprep system. RNA was sequenced on an Illumina HiSeq 4000. Analysis of these data was performed using the Ingenuity Pathway Analysis software (QIAGEN).

### UK Biobank

The UK biobank was accessed January 2019 and the data combined with the Hospital Episode data set (52). Subjects were excluded a priori if they took sleep altering medication or had obstructive sleep apnoea.

### Microarray Analysis

Geo2R (53) was used to analyse GSE47460 generated by the lung genome research consortium (54).

### Human PCLS

Precision cut lung slices were cut at 400μm on a vibrating microtome. TGFβ, GSK4112 or Vehicle (DMSO) treatments were performed each day with the slices being lysed after 4 days for qPCR analysis or 7 days for supernatant analysis. Additional methods can be found in the supplementary information

## Acknowledgements

We thank Leo Zeef and Andy Hayes of the Bioinformatics and Genomic Technologies Core Facilities at the University of Manchester for providing support with regard to the RNA-seq, Joe Takahashi for providing PER2∷Luc mice, the staff of the University of Manchester Histology, Bioimaging and Behavioral Sciences Facility for technical help. We would to thank the human study donors for their kind contribution.

## Funding

JB holds a MRC clinician scientist award (MR/L006499/1). DWR and AL both hold Wellcome Trust investigator awards (107851/Z/15/Z) and DWR is supported by a MRC programme grant (MR/P023576/1). JG has an Arthritis Research Career Development Fellowship (Reference 20629) and HD is supported by an Asthma UK Senior Clinical Academic Development Award (AUK-SCAD-2013-229). GK is supported by a MRC clinical research training fellowship (MR/N002024/1). The research of BH is supported by a Foundation Grant of the Canadian Institutes of Health Research (CIHR), and infrastructure Grants from the Canadian Foundation for Innovation (CFI) and Ontario Research Funds (ORF). N.C.H. is supported by a Wellcome Trust Senior Research Fellowship in Clinical Science (ref. 103749). LAB is supported by an MRC-MICA award (MR/R023026/1). AF is funded by the National Institute for Health Research Blood and Transplant Research Unit (NIHR BTRU) in Organ Donation and Transplantation at Newcastle University and in partnership with NHS Blood and Transplant (NHSBT). The Manchester Allergy, Respiratory and Thoracic Surgery Biobank is supported by the North West Lung Centre Charity and National Institute for Health Research Clinical Research Facility at Manchester University NHS Foundation Trust. This research was also supported by the NIHR Manchester Biomedical Research Centre.

## Notes

Conflict of Interest: The authors have declared that no conflict of interest exists

## References

1. Pariollaud M, et al. (2018) Circadian clock component REV-ERBalpha controls homeostatic regulation of pulmonary inflammation. The Journal of clinical investigation 128(6):2281–2296.

2. Gibbs J, et al. (2014) An epithelial circadian clock controls pulmonary inflammation and glucocorticoid action. Nat.Med. 20(8):919–926.

3. Cunningham PS, et al. (2019) Incidence of primary graft dysfunction after lung transplantation is altered by timing of allograft implantation. Thorax 74(4):413–416.

4. Pekovic-Vaughan V, et al. (2014) The circadian clock regulates rhythmic activation of the NRF2/glutathione-mediated antioxidant defense pathway to modulate pulmonary fibrosis. Genes & development 28(6):548–560.

5. Gibbs JE, et al. (2012) The nuclear receptor REV-ERBalpha mediates circadian regulation of innate immunity through selective regulation of inflammatory cytokines. Proc.Natl.Acad.Sci.U.S.A 109(2):582–587.

6. Gibbs JE, et al. (2009) Circadian timing in the lung; a specific role for bronchiolar epithelial cells. Endocrinology 150(1):268–276.

7. Jenkins RG, et al. (2017) An Official American Thoracic Society Workshop Report: Use of Animal Models for the Preclinical Assessment of Potential Therapies for Pulmonary Fibrosis. American journal of respiratory cell and molecular biology 56(5):667–679.

8. Cottin V, et al. (2018) Presentation, diagnosis and clinical course of the spectrum of progressive-fibrosing interstitial lung diseases. European respiratory review : an official journal of the European Respiratory Society 27(150).

9. Richeldi L, Collard HR, & Jones MG (2017) Idiopathic pulmonary fibrosis. Lancet (London, England) 389(10082):1941–1952.

10. Pardo A & Selman M (2016) Lung Fibroblasts, Aging, and Idiopathic Pulmonary Fibrosis. Annals of the American Thoracic Society 13 Suppl 5:S417–s421.

11. Hinz B (2012) Mechanical aspects of lung fibrosis: a spotlight on the myofibroblast. Proceedings of the American Thoracic Society 9(3):137–147.

12. Nureki SI, et al. (2018) Expression of mutant Sftpc in murine alveolar epithelia drives spontaneous lung fibrosis. The Journal of clinical investigation 128(9):4008–4024.

13. Moeller A, et al. (2009) Circulating fibrocytes are an indicator of poor prognosis in idiopathic pulmonary fibrosis. American journal of respiratory and critical care medicine 179(7):588–594.

14. Henderson NC, et al. (2013) Targeting of alphav integrin identifies a core molecular pathway that regulates fibrosis in several organs. Nature medicine 19(12):1617–1624.

15. Burgstaller G, et al. (2017) The instructive extracellular matrix of the lung: basic composition and alterations in chronic lung disease. The European respiratory journal 50(1).

16. Lodyga M, et al. (2019) Cadherin-11-mediated adhesion of macrophages to myofibroblasts establishes a profibrotic niche of active TGF-beta. Science signaling 12(564).

17. Takahashi JS (2017) Transcriptional architecture of the mammalian circadian clock. Nature reviews. Genetics 18(3):164–179.

18. Hatori M, et al. (2012) Time-restricted feeding without reducing caloric intake prevents metabolic diseases in mice fed a high-fat diet. Cell metabolism 15(6):848–860.

19. Archer SN, Schmidt C, Vandewalle G, & Dijk DJ (2018) Phenotyping of PER3 variants reveals widespread effects on circadian preference, sleep regulation, and health. Sleep medicine reviews 40:109–126.

20. Man K, Loudon A, & Chawla A (2016) Immunity around the clock. Science 354(6315):999–1003.

21. West AC, et al. (2017) Misalignment with the external light environment drives metabolic and cardiac dysfunction. Nature communications 8(1):417.

22. Kervezee L, Cermakian N, & Boivin DB (2019) Individual metabolomic signatures of circadian misalignment during simulated night shifts in humans. PLoS Biol 17(6):e3000303.

23. Bunger MK, et al. (2000) Mop3 is an essential component of the master circadian pacemaker in mammals. Cell 103(7):1009–1017.

24. Zhang Y, et al. (2015) GENE REGULATION. Discrete functions of nuclear receptor Rev-erbalpha couple metabolism to the clock. Science 348(6242):1488–1492.

25. Hinz B (2006) Masters and servants of the force: the role of matrix adhesions in myofibroblast force perception and transmission. European journal of cell biology 85(3-4):175–181.

26. Martin K, et al. (2016) PAK proteins and YAP-1 signalling downstream of integrin beta-1 in myofibroblasts promote liver fibrosis. Nature communications 7:12502.

27. Reed NI, et al. (2015) The alphavbeta1 integrin plays a critical in vivo role in tissue fibrosis. Sci Transl Med 7(288):288ra279.

28. Gerber EE, et al. (2013) Integrin-modulating therapy prevents fibrosis and autoimmunity in mouse models of scleroderma. Nature 503(7474):126–130.

29. Collins R (2012) What makes UK Biobank special? Lancet (London, England) 379(9822):1173–1174.

30. Shan Z, et al. (2015) Sleep duration and risk of type 2 diabetes: a meta-analysis of prospective studies. Diabetes care 38(3):529–537.

31. Knutson KL & von Schantz M (2018) Associations between chronotype, morbidity and mortality in the UK Biobank cohort. Chronobiology international 35(8):1045–1053.

32. Vetter C, et al. (2018) Night Shift Work, Genetic Risk, and Type 2 Diabetes in the UK Biobank. Diabetes care 41(4):762–769.

33. Bauer Y, et al. (2015) A novel genomic signature with translational significance for human idiopathic pulmonary fibrosis. American journal of respiratory cell and molecular biology 52(2):217–231.

34. Meng QJ, et al. (2008) Ligand modulation of REV-ERBalpha function resets the peripheral circadian clock in a phasic manner. J Cell Sci. 121(Pt 21):3629–3635.

35. Bonniaud P, et al. (2005) Progressive transforming growth factor beta1-induced lung fibrosis is blocked by an orally active ALK5 kinase inhibitor. American journal of respiratory and critical care medicine 171(8):889–898.

36. Gerber A, et al. (2013) Blood-borne circadian signal stimulates daily oscillations in actin dynamics and SRF activity. Cell 152(3):492–503.

37. Balsalobre A, Damiola F, & Schibler U (1998) A serum shock induces circadian gene expression in mammalian tissue culture cells. Cell 93(6):929–937.

38. Hoyle NP, et al. (2017) Circadian actin dynamics drive rhythmic fibroblast mobilization during wound healing. Sci Transl Med 9(415).

39. Kowalska E, et al. (2013) NONO couples the circadian clock to the cell cycle. Proceedings of the National Academy of Sciences of the United States of America 110(5):1592–1599.

40. Jones SE, et al. (2019) Genome-wide association analyses of chronotype in 697,828 individuals provides insights into circadian rhythms. Nature communications 10(1):343.

41. Lane JM, et al. (2019) Biological and clinical insights from genetics of insomnia symptoms. Nature genetics 51(3):387–393.

42. Thorn JC, et al. (2016) Validation of the Hospital Episode Statistics Outpatient Dataset in England. PharmacoEconomics 34(2):161–168.

43. Raghu G, et al. (2011) An official ATS/ERS/JRS/ALAT statement: idiopathic pulmonary fibrosis: evidence-based guidelines for diagnosis and management. American journal of respiratory and critical care medicine 183(6):788–824.

44. Grandner MA & Drummond SP (2007) Who are the long sleepers? Towards an understanding of the mortality relationship. Sleep medicine reviews 11(5):341–360.

45. Jike M, Itani O, Watanabe N, Buysse DJ, & Kaneita Y (2018) Long sleep duration and health outcomes: A systematic review, meta-analysis and meta-regression. Sleep medicine reviews 39:25–36.

46. Trump RP, et al. (2013) Optimized chemical probes for REV-ERBalpha. J.Med.Chem. 56(11):4729–4737.

47. Solt LA, et al. (2012) Regulation of circadian behaviour and metabolism by synthetic REV-ERB agonists. Nature 485(7396):62–68.

48. Dierickx P, et al. (2019) SR9009 has REV-ERB-independent effects on cell proliferation and metabolism. Proceedings of the National Academy of Sciences of the United States of America 116(25):12147–12152.

49. Carrington R, Jordan S, Pitchford SC, & Page CP (2018) Use of animal models in IPF research. Pulmonary pharmacology & therapeutics 51:73–78.

50. Yoo SH, et al. (2004) PERIOD2∷LUCIFERASE real-time reporting of circadian dynamics reveals persistent circadian oscillations in mouse peripheral tissues. Proceedings of the National Academy of Sciences of the United States of America 101(15):5339–5346.

51. Smith-Clerc J & Hinz B (2010) Immunofluorescence detection of the cytoskeleton and extracellular matrix in tissue and cultured cells. Methods in molecular biology (Clifton, N.J.) 611:43–57.

52. Herbert A, Wijlaars L, Zylbersztejn A, Cromwell D, & Hardelid P (2017) Data Resource Profile: Hospital Episode Statistics Admitted Patient Care (HES APC). International journal of epidemiology 46(4):1093–1093i.

53. Barrett T, et al. (2013) NCBI GEO: archive for functional genomics data sets--update. Nucleic acids research 41(Database issue):D991–995.

54. Yu G, et al. (2018) Thyroid hormone inhibits lung fibrosis in mice by improving epithelial mitochondrial function. Nature medicine 24(1):39–49.

