## Supplemental Information for "The circadian clock protein REVERBα inhibits pulmonary fibrosis development"

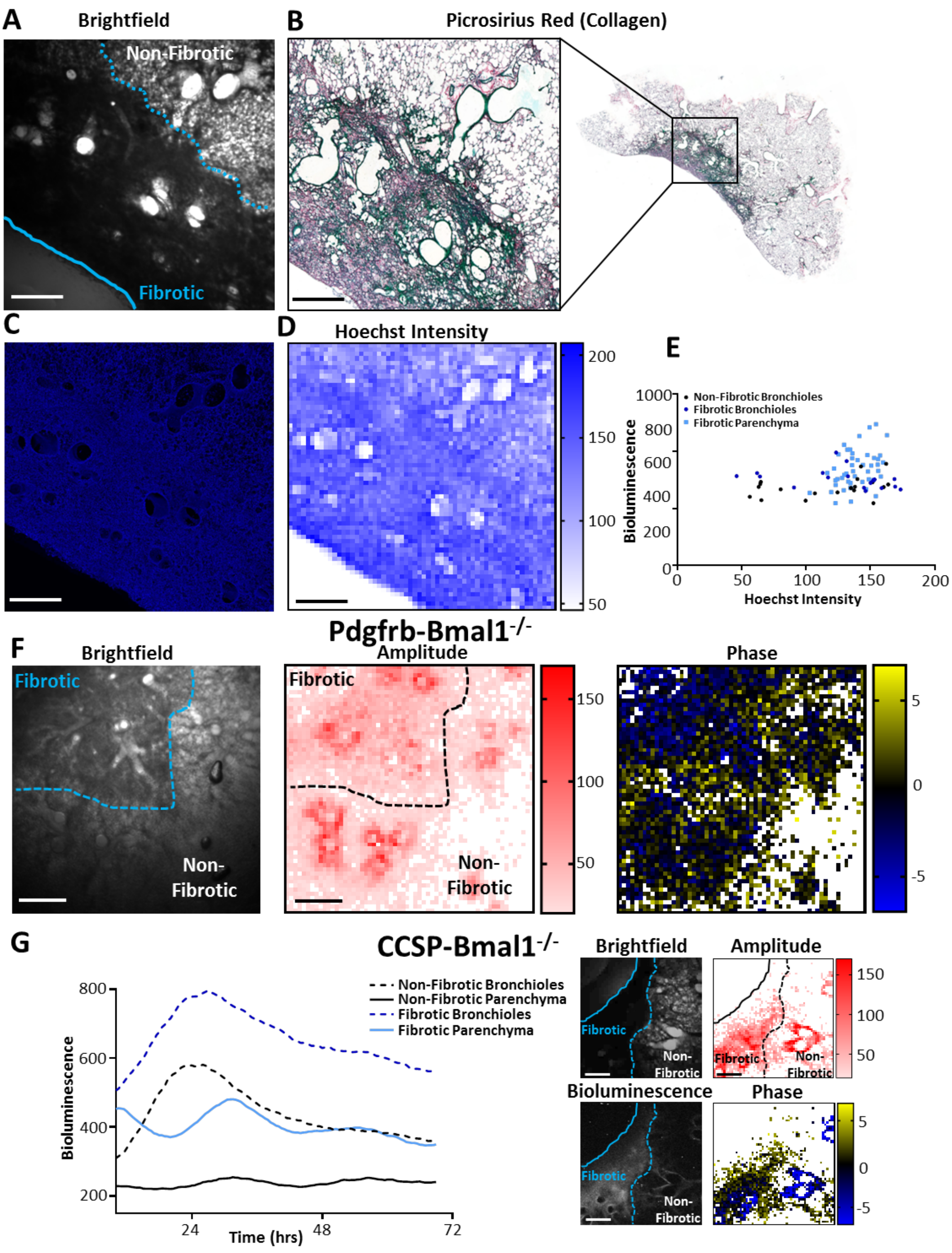


**Figure S1 - Analysis of fibrotic mPER2::luc precision cut lung slices.**
**A)** Brightfield image of the precision cut lung slice used in Fig. 1A **B)** Histological section of the precision cut lung slice used in Fig. 1A stained with picrosirius red **C)** Immunofluorescence of the precision cut lung slice used in Fig1A following Hoechst staining

**D)** Heatmap (1500µm$^2$) of Hoechst intensity in the same precision cut lung slice **E)** Hoechst intensity was plotted against bioluminescence for the precision cut lung slice in different anatomical structures, no significant correlations were observed **F)** Brightfield image**,** Amplitude and Phase heatmaps (1500µm$^2$) of PER2::luc oscillations in the precision cut lung slice from the Pdgfrb-BMAL1$^{-/-}$ mouse; Scale bars, 500µm (data is representative of three separate experiments) **G)** Representative bioluminescent image along with bioluminescent intensity, Brightfield image**,** Amplitude and Phase heatmaps (1500µm$^2$) of PER2::luc oscillations in the precision cut lung slice from the CCSP-BMAL1$^{-/-}$ mouse; (Scale bars, 500µm, data is representative of three separate experiments).

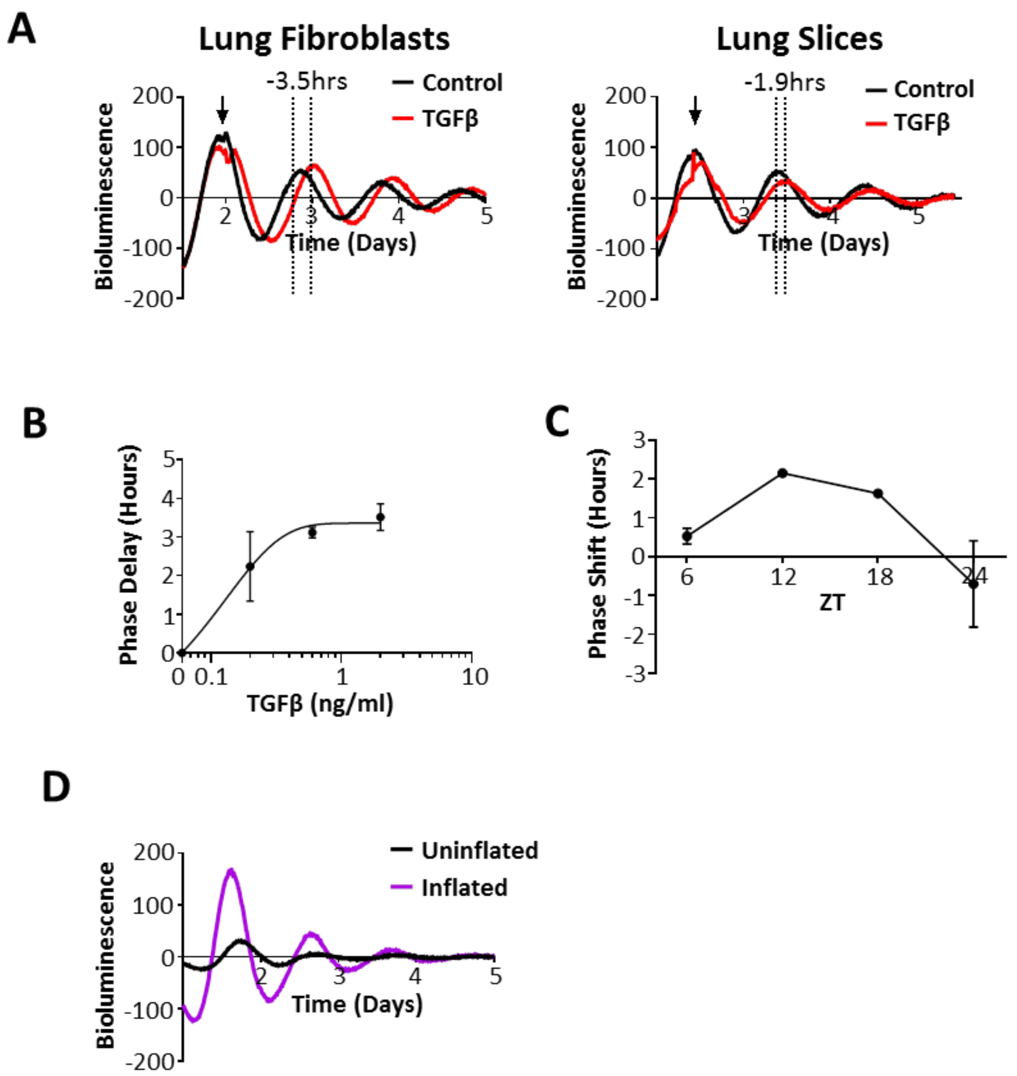


**Figure S2 - Effect of TGFβ and lung inflation on PER2 rhythmicity in primary murine fibroblasts and lung slices**

**A)** Bioluminescence was plotted against time for fibroblasts and precision cut lung slices from mPER2::luc mice. TGFβ (2ng/ml) was added at the time-point indicated by the arrow (data from one representative experiment but is representative of three experiments) with the subsequent peaks shown by the dotted lines **B)** Dose response curve of mPER2::luc primary lung fibroblasts following treatment with TGFβ (mean±SEM, data is plotted from three experiments) **C)** Phase response curve in mPER2::luc precision cut lung slices following TGFβ (2ng/ml) stimulation (mean±SEM, 4 slices per time point all from separate animals) **D)** Lungs from mPER2::luc mice where inflated or left uninflated. Bioluminescence was then plotted against time (representative data from one animal, (n=3 animals)).

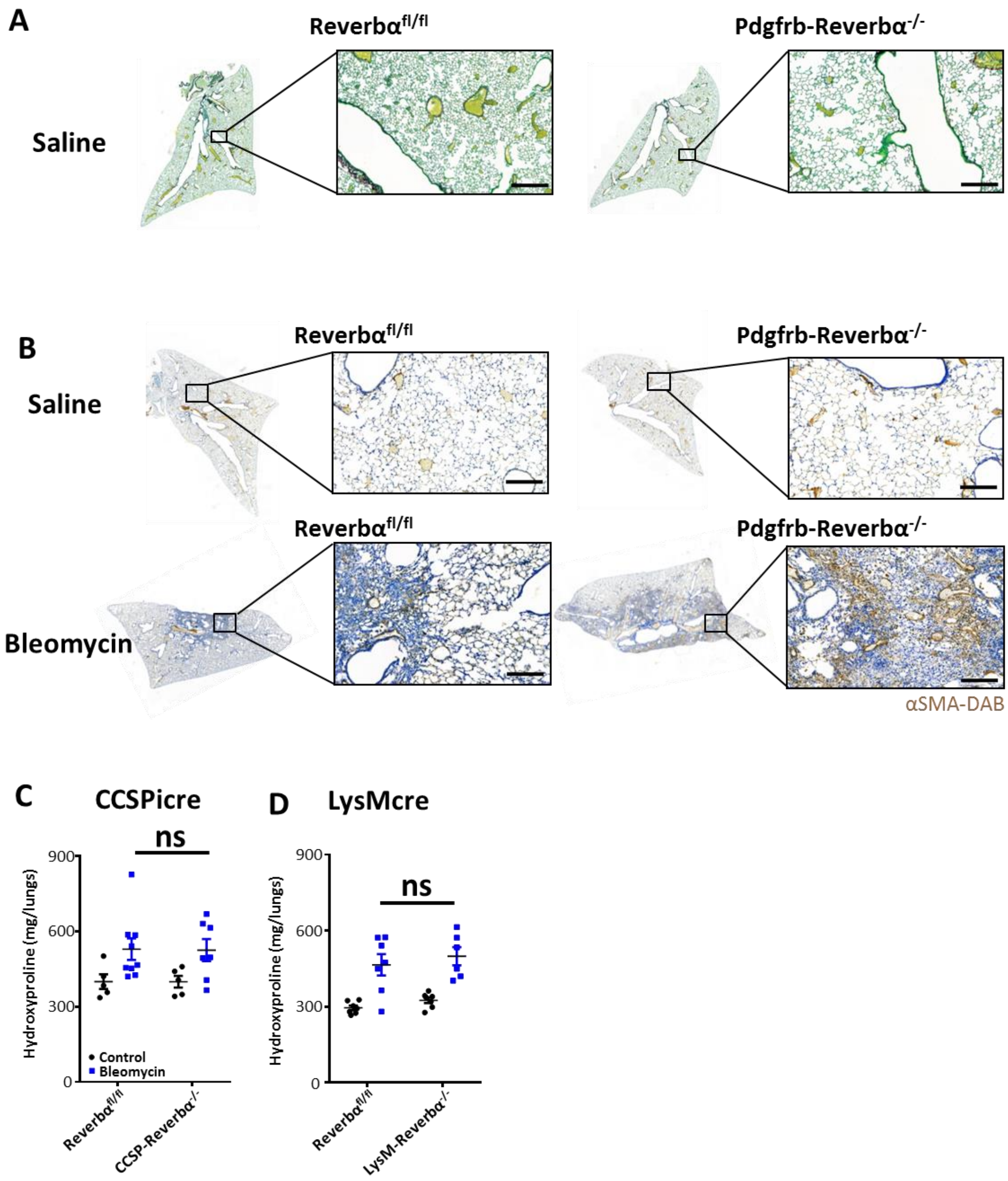


**Figure S3 - Myofibroblasts are increased in the Pdgfrb-Reverbα$^{-/-}$ mouse, whereas conditional modulation of REVERBα in club cells or myeloid cells fails to alter the pulmonary fibrotic response to bleomycin.**

**A)** Histological image stained with Picrosirius red of representative tissue sections from saline treated mice (3 animals per genotype); scale bar, 200µm. **B)** Representative lung images after immunohistochemistry (anti-αSMA, 3,3′-Diaminobenzidine (DAB)) 28 days after intra-tracheal bleomycin or saline challenge (3 animals treated with saline and 4-5 animals treated with bleomycin per genotype; scale bar, 200µm). The bleomycin inset image has been shown in Fig. 2D **C)** Hydroxyproline measurements from CCSPicre-Reverbα$^{-/-}$ vs littermate controls 28 days after intra-tracheal bleomycin (2U/kg) challenge (mean±SEM; n=5 saline and 7-9 bleomycin per genotype) **D)** Hydroxyproline measurements from LysM-Reverbα$^{-/-}$ mice vs littermate controls 28 days after intra-tracheal bleomycin (3U/kg) challenge (mean±SEM; n=5 saline and 6-7 bleomycin per genotype).

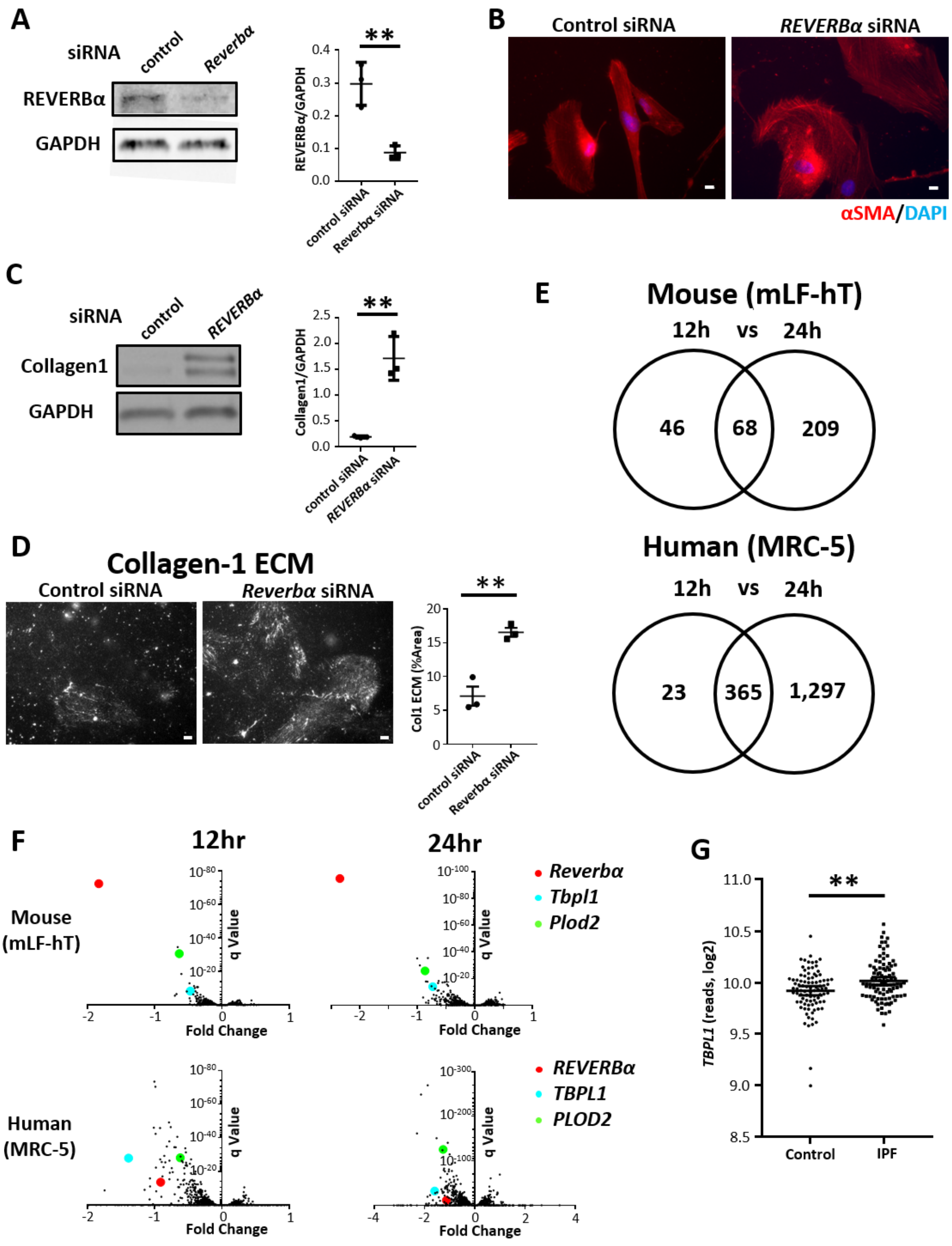


**Figure S4 – RNA-seq reveals REVERBα's effect on TBPL1 expression which is also altered in patients with idiopathic pulmonary fibrosis**

**A)** Immunoblot for REVERBα following transfection of control (non-targeting) or Reverbα (*nr1d1*) siRNA in mLF-hT cells. (representative immunoblot from n=3 transfections, **=P<0.01 Student's t-test) **B)** Immunofluorescent staining for the myofibroblast marker αSMA after Control (non-targeting) or *REVERBα* siRNA knockdown in MRC-5 cells; scale bar,

10µm (representative image from 3 separate transfections) **C)** Immunoblot for Collagen-1 protein following transfection of control (non-targeting) or *REVERBα* siRNA knockdown in MRC-5 cells (**=P<0.01 student t test, n=3 transfections) **D)** Immunofluorescent staining and quantification of Collagen-1 ECM deposition following transfection of control (non-targeting) or *Reverbα* siRNA knockdown in mLF-hT cells. (n=3 transfections, representative image shown, **=P<0.01 Student t-test); scale bar, 50µm **E)** Venn diagrams (12hrs Vs 24hrs) showing genes differentially (q<0.05) regulated using RNA-seq analysis in MRC-5 and mLF-hT cells following siRNA silencing of REVERBα (n=3 transfections per sampling point) **F)** Volcano plots (q value vs fold change) for each of the four RNA-seq experimental conditions from which the mean plot is shown in Fig. 3C (n=3 transfections per sampling point) **G)** Changes in *TBPL1* gene expression in idiopathic pulmonary fibrosis compared to control subjects from a previously published genome array (GSE 47460) (fold change ± 95% confidence interval, n= 90 controls and 98 patients with IPF) **=P<0.01 Student's t-test.

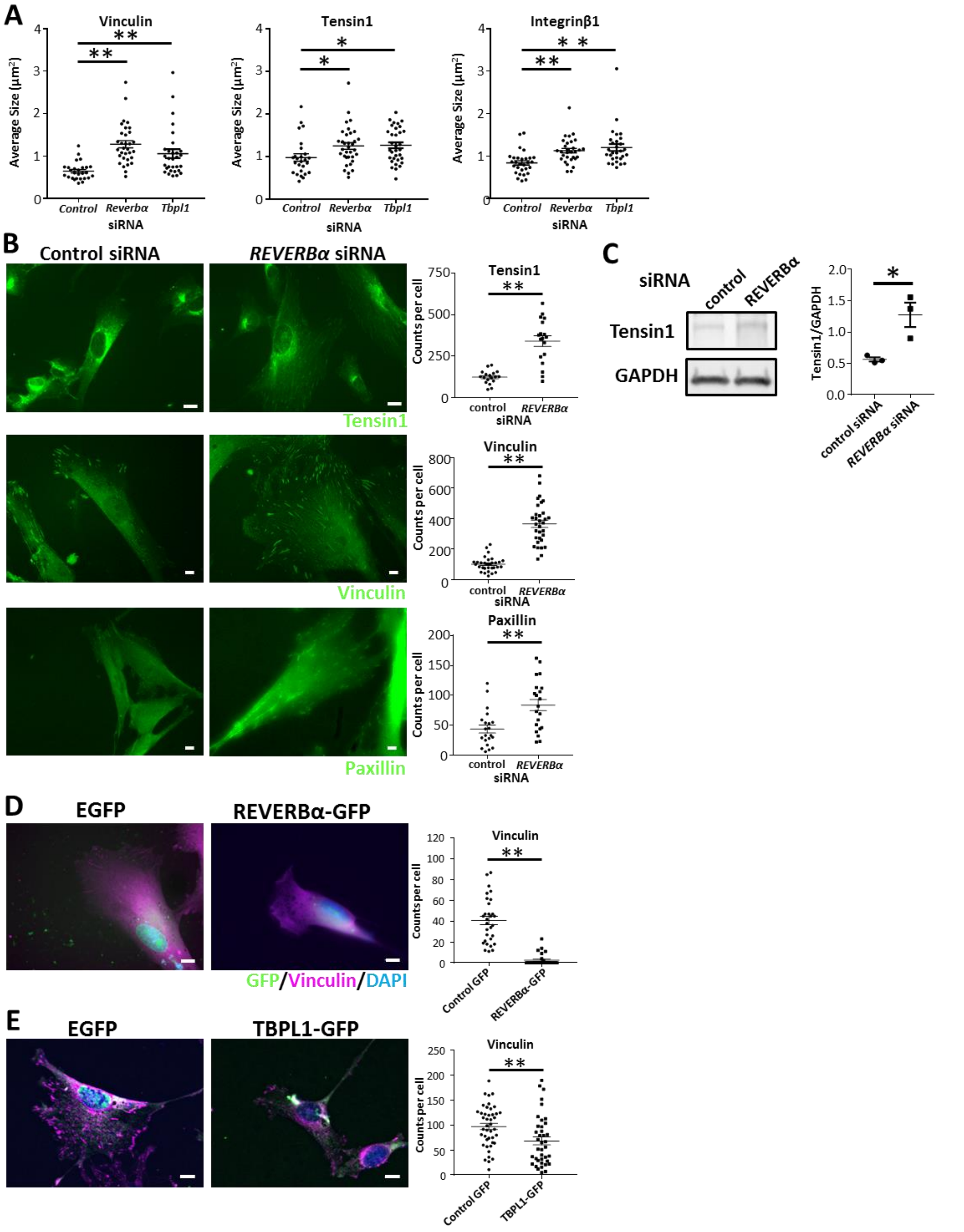


**Figure S5 – Characterization of cell changes in mLF-hT and Mrc5 cells following either REVERBα knockdown or REVERBα/ TBPL1 overexpression.**

**A)** Quantification of, vinculin, tensin1 and integrinβ1 average cluster size following siRNA knockdown of *Reverbα* or *Tbpl1* compared to control (non-targeting) siRNA in mLF-hT cells (****=**p<0.01 ANOVA •= individual cells from 3 transfections; scale bar, 10µm **B)**

Representative images and quantification of tensin-1, vinculin and paxillin immunofluorescence in MRC-5 cells following *REVERBα* or Control (non-targeting) siRNA knockdown. (****=**p<0.01 Student's t-test •= individual cells from 3 transfections; scale bar, 10µm) **C)** Representative immunoblot along with quantification of tensin1 following *REVERBα* or Control (non-targeting) siRNA knockdown in MRC-5 cells. (representative immunoblot from 3 separate transfections *=p<0.05 Student's t-test) **D)** Representative immunofluorescence images following transfection with REVERBα-GFP or eGFP along with quantification of focal adhesions (vinculin) in MRC5 cells. Cells were stained for GFP, Vinculin and nuclei (DAPI) (**=p<0.01 student's t-test, •= individual cells from 3 separate transfections) **E)** Representative immunofluorescence images following transfection with TBPL1-GFP or eGFP along with quantification of focal adhesions (vinculin) in mLF-hT cells. Cells were stained for GFP, Vinculin and nuclei (DAPI) (****=**p<0.01 student's t-test, •= individual cells from 3 separate transfections).

**Supplementary Videos:**

**Video S1: Per2 bioluminescence in WT mice after bleomycin** Video showing bioluminescence over time in a precision cut lung slice in a WT mouse on a mPER2::luc background. The mouse was treated with intra-tracheal bleomycin 14 days before the lung slice was harvested.

**Video S2: Per2 bioluminescence in CCSPicre-Bmal1$^{-/-}$ mice after bleomycin** Video showing bioluminescence over time in a precision cut lung slice in a CCSPicre-Bmal1$^{-/-}$ on a mPER2::luc background. The mouse was treated with intra-tracheal bleomycin 14 days before the lung slice was harvested.

**Video S3: Per2 bioluminescence in Pdgfrb-Bmal1$^{-/-}$ mice after bleomycin** Video showing bioluminescence over time in a precision cut lung slice in a Pdgfrb-Bmal1$^{-/-}$ on a mPER2::luc background. The mouse was treated with intra-tracheal bleomycin 14 days before the lung slice was harvested.

### Supplementary Tables:

| Chronotype | Number of cases | Sample size | Odds Ratio | P value | 95% Confidence Interval | |
|---|---|---|---|---|---|---|
| Morning | 753 | 276,533 | 1 | - | - | - |
| Evening | 493 | 165,515 | 1.030 | 0.121 | 0.992 | 1.070 |

**Table S1: Unadjusted Association between Chronotype and Pulmonary Fibrosis:** Odds ratios were calculated for the association between evening chronotype and pulmonary fibrosis using binary logistic regression using volunteers from the UK biobank

| Variable | Number of cases | Sample size | Odds Ratio | P value | 95% Confidence Interval | |
|---|---|---|---|---|---|---|
| Morning Chronotype | 753 | 276,533 | 1 | - | 1 | 1 |
| Evening Chronotype | 493 | 165,515 | 1.040 | 0.047 | 1.001 | 1.080 |
| Age | - | - | 1.103 | <0.001 | 1.093 | 1.113 |
| Male Sex | - | - | 1.387 | <0.001 | 1.556 | 1.237 |
| BMI | - | - | 1.035 | <0.001 | 1.023 | 1.047 |
| Smoking | - | - | 2.128 | <0.001 | 1.896 | 2.388 |

**Table S2: Adjusted association between Chronotype and Pulmonary Fibrosis:** Odds ratios were calculated for the association between evening chronotype and pulmonary fibrosis using binary logistic regression after adjusting for confounders (sex, age, BMI and smoking)

| Shift work | Number of cases | Sample size | Odds Ratio | P value | 95% Confidence Interval | |
|---|---|---|---|---|---|---|
| None | 335 | 236,190 | 1 | - | 1 | 1 |
| Yes | 90 | 49,370 | 1.286 | 0.034 | 1.019 | 1.623 |

**Table S3: Unadjusted association between Shift work and Pulmonary Fibrosis:** Odds ratios were calculated for the association between shift work and pulmonary fibrosis using binary logistic regression

| Variable | Number of cases | Sample size | Odds Ratio | P value | 95% Confidence Interval | |
|---|---|---|---|---|---|---|
| No Shift work | 335 | 236,190 | 1 | - | 1 | 1 |
| Shift Work | 90 | 49,370 | 1.353 | 0.012 | 1.069 | 1.711 |
| Age | - | - | 1.106 | <0.001 | 1.090 | 1.122 |
| Sex | - | - | 1.238 | 0.032 | 1.017 | 1.506 |
| BMI | - | - | 1.031 | 0.003 | 1.010 | 1.052 |
| Smoking | - | - | 1.622 | <0.001 | 1.336 | 1.968 |

**Table S4: Adjusted association between Shift work and Pulmonary Fibrosis:** Odds ratios were calculated for the association between shift work and pulmonary fibrosis using binary logistic regression after adjusting for confounders (sex, age, BMI and smoking)

| Sleep duration (h) | Number of cases | Sample size | Odds Ratio | P value | 95% Confidence Interval | |
|---|---|---|---|---|---|---|
| ≤4 | 24 | 5,585 | 1.974 | <0.001 | 1.307 | 2.981 |
| 5 | 94 | 21,760 | 1.985 | <0.001 | 1.586 | 2.483 |
| 6 | 287 | 95,060 | 1.385 | <0.001 | 1.192 | 1.610 |
| 7 | 418 | 191,619 | 1 | - | 1 | 1 |
| 8 | 404 | 143,771 | 1.289 | <0.001 | 1.124 | 1.478 |
| 9 | 133 | 28,982 | 2.109 | <0.001 | 1.734 | 2.564 |
| 10 | 43 | 7,027 | 2.816 | <0.001 | 2.056 | 3.858 |
| ≥11 | 18 | 2,107 | 3.941 | <0.001 | 2.454 | 6.330 |

**Table S5: Unadjusted association between Sleep duration and Pulmonary Fibrosis.** Odds ratios were calculated for the association between shift work and pulmonary fibrosis using binary logistic regression

| Variable | Number of cases | Sample size | Odds Ratio | P value | 95% Confidence Interval | |
|---|---|---|---|---|---|---|
| ≤4h sleep | 24 | 5,585 | 1.836 | 0.004 | 1.214 | 2.776 |
| 5h sleep | 94 | 21,760 | 1.808 | <0.001 | 1.444 | 2.265 |
| 6h sleep | 287 | 95,060 | 1.314 | <0.001 | 1.129 | 1.528 |
| 7h sleep | 418 | 191,619 | 1 | - | 1 | 1 |
| 8h sleep | 404 | 143,771 | 1.134 | 0.074 | 0.988 | 1.302 |
| 9h sleep | 133 | 28,982 | 1.542 | <0.001 | 1.265 | 1.880 |
| 10h sleep | 43 | 7,027 | 1.947 | <0.001 | 1.414 | 2.682 |
| ≥11h sleep | 18 | 2,107 | 3.149 | <0.001 | 1.955 | 5.071 |
| Age | - | - | 1.100 | <0.001 | 1.091 | 1.110 |
| Male Sex | - | - | 1.421 | <0.001 | 1.276 | 1.583 |
| BMI | - | - | 1.029 | <0.001 | 1.017 | 1.040 |
| Smoking | - | - | 2.059 | <0.001 | 1.849 | 2.292 |

**Table S6: Adjusted association between Sleep duration and Pulmonary Fibrosis:** Odds ratios were calculated for the association between shift work and pulmonary fibrosis using binary logistic regression after adjusting for confounders (sex, age, BMI and smoking)

**Supplementary Materials and Methods:**

**UK Biobank and HES Analysis**

The UK biobank recruited 502,620 individuals in the UK population between 2006 and 2010. Participants filled in a questionnaire regarding lifestyle, medical conditions along with their demographic information. The questionnaire included questions on chronotype, shift work and sleep duration and has been validated in other studies (1). The medical information collected by the UK biobank was supplemented by the English hospital episodes data set (HES) (2) accessed January 2019. Subjects were excluded *a priori* if they took sleep altering medication (3) or had obstructive sleep apnoea. Pulmonary fibrosis was defined using UK biobank codes 1121,1122 and 1115 combined with ICD 10 codes J84.1 and J84.9 in the HES dataset.

Shift work was defined using a computer administered questionnaire were participants who worked were asked if they worked shifts. Possible answers were "never/rarely", "sometimes", "usually", "always", "prefer not to answer" and "do not know". If the participant checked either of the last two options, they were excluded from shift work analysis. Patients were defined as non-shift workers if they ticked the box "never/rarely". Chronotype was defined in a similar manner using the same touchscreen questionnaire based on the morningness-eveningness questionnaire. Due to the low incidence of pulmonary fibrosis the two answers signifying a morning chronotype were pooled and the same was done for the evening chronotype. This single item has been shown to correlate with sleep timing and dim-light melatonin onset (4). Sleep duration was obtained from the same touchscreen questionnaire where subjects in response to the question "About how many hours sleep do you get in every 24 hours? (please include naps)?"(5) with the response being recorded in hours.

**Microarray analysis**

Geo2R (6) was used to analyse GSE47460 generated by the lung genome research consortium (7) who performed microarray analysis on lung tissue from patients with pulmonary fibrosis as well as control subjects. Patients were classified as having clinically significant pulmonary fibrosis based on reduced lung function (FVC<80%predicted) and a pathological diagnosis of UIP.

#### *In vivo* Bleomycin model of pulmonary fibrosis

Male mice (aged 10–14 weeks) were dosed intratracheally with bleomycin (Sigma) (50µL) or vehicle (saline) under anaesthesia (isoflurane). Bleomycin and vehicle treated animals were analysed 28 days after exposure unless stated otherwise. Hydroxyproline was measured on snap frozen lungs, histological analysis was performed on lungs inflated with 4% paraformaldehyde (PFA) that were then left to fix overnight. For precision cut lung slices, lungs were harvested 14-21 days after bleomycin.

#### Hydroxyproline measurement of tissue collagen

Hydroxyproline was measured using a colourimetric assay as previously described (8). Briefly, whole lungs were dissected, weighed and homogenised in a mortar and pestle under liquid nitrogen. ~50mg of tissue was dried (80°C for 1 hour) and then hydrolysed in 6N HCl at 100°C for 20 hours. Samples were then oxidised in Citrate-Acetate Buffer with Chloramine T allowing colourimetric determination of hydroxyproline concentration with p-diamethylaminobenzaldehyde in Isopropanol/perchloric acid.

#### Bioluminescence Microscopy

Precision-cut organotypic lung slices (PCLS) were prepared as described before (9). Briefly, the trachea was cannulated with flexible polyethylene tubing of 0.58 mm in diameter. The lungs were inflated with 2 ml of freshly prepared agarose (2% w/v in HBSS media at 38°C) or until the accessory lobe was fully inflated. The trachea was tied off with 2/0 nylon and the lungs were then placed on ice until the agarose had set (approx. 15mins). The lungs were then fixed onto the stage of a vibrating microtome (Campden instruments integraslice 7550mm) and sectioned at 250µm at 4°C.

After sectioning the slices underwent sequential washes of DMEM to remove agarose and were then placed on a cell culture insert (MilliCell cell culture inserts 0.04µm pores) in luciferin containing culture media (phenol red-free DMEM with 3.5 mg/mL glucose, 25 U/mL penicillin, 25 µg/mL streptomycin, 0.1 mM luciferin, 10 mM HEPES-NaOH)(9). Brightfield imaging was captured on a Zeiss Axiovert 100 with a Plan Achromat 5× 0.16 NA objective equipped with an XL incubator to maintain the cells at 37°C, 5% $CO_2$. Bioluminescence images were obtained using a 2.5x objective (Zeiss) and captured using a cooled Andor iXon Ultra camera over a 30 minute integration period acquired using Micro-Manager software (Version 1.4).

Captured images were analysed using MATLAB R2018a to define amplitude and phase of oscillations. In brief a grid was placed across the image where each square represented 1500μm$^2$. The same analysis grid was placed on subsequent images captured 30 minutes apart allowing phase and amplitude to be calculated for each defined area. Spikes over time were removed via Hampel outlier identification (10) in order to reduce the influence of random photons and then baseline subtraction was performed for each square using a 24 hour moving average. The acrophase was then calculated by locating the local maxima of the time series. The prominence and relative timing of the acrophase were used to calculate respectively amplitude (in arbitrary units) and phase differences (in hours) per square. Amplitude and phase was then plotted into a heatmap where the location of each squares corresponds with its morphological position in the original lung slice.

**Hoechst staining**

After bioluminescent imaging the tissue was fixed in 4% PFA for 2 hours and stored in 70% ethanol at 4°C until Hoechst staining. PCLS were then incubated in PBS with 2 μM Hoechst 33342 for 15 minutes. Staining solution was removed and replaced by PBS. Nuclei were imaged using a Leica TCS SP8 AOBS upright confocal microscope to obtain a 3D optical stack of the whole lung slice. The image presented in the results section is a maximum intensity projection of this 3D stack. Using morphological landmarks, the field of view of this image was matched with the field of view of the previous bioluminescence recording.

**Lumicycle Bioluminescence Analysis**

Precision cut lung slices and primary lung fibroblasts were also analysed with a lumicycle bioluminescence recorder after being placed on cell culture inserts in a 35mm dishes containing media with luciferin as previously described (11). To investigate the effects of lung inflation, lungs were either inflated with 2% agarose solution and then set at 4°C or kept uninflated at 4°C. Sections of lung were then cut and placed in the lumicycle.

**Histology and Immunohistochemistry**

Paraformaldehyde fixed whole lungs or precision cut lung slices were paraffin embedded and cut sectioned at 5µm. To visualise collagen, sections were dewaxed, rehydrated and stained with Picrosirius Red/Fast Green. For αSMA IHC dewaxed and rehydrated sections were processed for antigen retrieval followed by blocking for endogenous peroxidase with BLOXALL and Mouse IgGs (Vector Laboratories). Sections were then incubated with anti-αSMA antibody overnight prior to detection and staining (MOM ImmPRESS Peroxidase Polymer Kit (Vector Laboratories) and DAB (3'3'-diaminobenzidine)). Control immunohistochemistry was also performed on sections without the addition of either primary or secondary antibodies to confirm specificity. αSMA IHC was graded (0-4) by scorers blinded to the sample details based on intensity of the αSMA stain and the percentage of tissue affected.

**Generation of primary mouse lung fibroblasts**

Whole lungs were dissected to ~2mm$^3$ pieces and washed in Hank's Balanced Salt Solution (HBSS, Sigma-Aldrich). This tissue was then Collagenase (1.5mg/ml in HBSS, Sigma-Aldrich) digested at 37°C for 2 hours. After Collagenase incubation the homogenate was passed through a 40µm cell strainer, washed in HBSS and finally re-suspended in culture media (DMEM +10% FBS) for fibroblast cell expansion.

**Cell transfections**

Twenty-four hours post-seeding (imFLs, 1.5 X $10^3$ cells/cm$^2$; Mrc5 0.5 X $10^4$ cells/cm$^2$) siRNA (15ng siRNA/cm$^2$; mouse *Nr1d1* (ON-TARGETplus SMARTpool #L-051721-00, Dharmacon), mouse *Tbpl1* (ON-TARGETplus SMARTpool #L-058115-01, Dharmacon), mouse *Itgb1* (ON-TARGETplus SMARTpool #L-040783-01, Dharmacon), human *NR1D1* (Silencer Select #s18387, Ambion), human *TBPL1* (ON-TARGETplus SMARTpool #L-017254-00, Dharmacon) and Non-targeting (ON-TARGETplus Control Pool #D-001810-10, Dharmacon)) transfections were performed using DharmaFECT 1 (Dharmacon) reagent.

To create a Rev-erbα-GFP plasmid Phsuion HF Taq was used to clone Rev-erbα using the following primers CGGCGGATCCATGACGACCCTGGACT (forward) and GCGCGGCCGCTCACTGGGCGTCCACCCG (reverse) on cDNA extracted from A549 cells according to manufacturer's instructions. The PCR product was then inserted into pGEM T easy Vector (Promega) and then sub-cloned into a pcDNA3-gfp vector (Addgene). The subsequent plasmid was then sequenced. Tbpl1-GFP (Origene) and control EGFP (Addgene) transfections were performed 24hours post-seeding using Fugene HD (2µg plasmid/10µl Fugene HD, Promega) as per the manufacturer's instructions.

**Immunofluorescence**

For αSMA staining cells in 35mm dishes were fixed in 4% PFA/0.2% Triton X, followed by ice cold methanol fixation (αSMA). For focal adhesions proteins cells were exposed to ice-cold cytoskeleton buffer (12) for 10 minutes followed by 4% PFA fixation for a further 10 minutes. Cells were then incubated with primary antibodies (indicated in the relevant figures) for 1 hour at room temperature and then washed prior to incubation with a species specific fluorescent secondary antibody (Alexa Flour) for one hour. Following this cell nuclei were then stained with DAPI before dishes were washed, mounted (Prolong Diamond) and cover slipped.

For ECM analysis cells were lysed in 20mM $NH_4OH$, 0.5% Triton X-100 in PBS, the DNA was digested using DNAse1 (Roche) and the remaining protein was stained with collagen-1 antibody.

Fiji (13) was used to analyse the immunofluorescence images. αSMA and collagen-1 were quantified using a macro which measured intensity above a threshold which was set the same for all experimental conditions. Focal adhesions were quantified via subtracting the background (rolling ball 50), then a fft band pass filter was applied allowing a threshold to be applied which identified the focal adhesions. All samples were coded and images were quantified by a user blinded to the cell transfection conditions.

**Antibodies**

The following antibodies were used throughout the paper; αSMA (mouse, Leica, NCL-L-SMA), Collagen Type 1 (goat, Southern Biotech, 1310-01), pro-Collagen1 (rabbit, R1038, Acris), Integrinβ1(Mouse) (Rat, BD Pharmingen, 550531), Integrinβ1(Human/active) (Mouse, Millipore, MAB2079Z), Vinculin (mouse, Sigma-Aldrich, V9131), Tensin1 (rabbit, Sigma-Aldrich, SAB4200283), Paxillin (mouse, 610569, BD Biosciences), REV-ERBα (mouse monoclonal, as previously described (14)), TBPL1 (rabbit, Proteintech, 12258-1-AP), GFP (rabbit, Abcam, ab290), GAPDH (rabbit, Proteintech, 10494-1-AP). Alexa Fluor fluorescent secondary antibodies (anti-rabbit-488 (A11008); anti-rat-568 (A11077) and anti-mouse-647 (A21235)) were purchased from Invitrogen.

**RNA-seq**

siRNA transfected mLF-hT and Mrc5 cells were lysed and RNA was extracted using the ReliaPrep RNA miniprep system. RNA was sequenced on an Illumina HiSeq 4000. Analysis of these data was performed using the Ingenuity Pathway Analysis software (QIAGEN).

**Human lung slice and fibroblast culture**

Human lung tissue was collected from patients undergoing carcinoma resection (control) or for fibrotic tissue from human explants after transplantation. Tissue which the pathologist deemed suitable for research was then inflated with 2-3% low boiling point agarose, with the agarose being allowed to set at 4°C. Precision cut lung slices were then cut at 400µm on a vibrating microtome and cultured in DMEM media. TGFβ (2ng/ml), GSK4112 (10µM) or Vehicle (DMSO) treatments were then performed each day with the slices being lysed after 4 days for qPCR analysis or 7 days for supernatant analysis.

To prepare a lung fibroblast culture, fibrotic and non-fibrotic tissue was dissected to ~2mm$^3$ pieces and placed in culture media in a 15cm$^2$ dish scored to create points of adhesion. Following 2-4 weeks, outgrown fibroblasts were trypsinised and passaged for further culture. For treatments, cells were plated in 24 well plates (1 X $10^5$ cells/well) and TGFβ (2ng/ml), GSK4112 (10µM) or Vehicle (DMSO) was added in serum-free culture media for 24 hours after which cells were lysed for qPCR analysis.

**ELISA**

Collagen 1a1 concentration in cell culture media from PCS were assessed using a DuoSet® kit (R&D Systems) as per manufacturer's instructions.

**Western immunoblotting**

For whole cell analyses, protein lysates were generated by lysis in RIPA buffer (Cell Signalling) containing Protease Inhibitor Cocktail (Roche) and 1 mM PMSF. Extracellular matrix protein was prepared by adding 4X NuPAGE LDS Sample Buffer (Novex) directly to the tissue culture plastic and scraping to collect the ECM after the prior removal of the cells with extraction buffer (20mM $NH_4OH$. 0.5% Triton X-100 in PBS). SDS-PAGE was performed using the mini-PROTEAN system (Bio-Rad) and protein levels were determined by chemiluminescent detection of target proteins on nitrocellulose membranes.

**qPCR**

RNA was extracted using the ReliaPrep RNA miniprep system (Promega) prior to the generation of cDNA (RNA-to-cDNA kit, Applied Biosystems). qPCR was performed using PowerUp SYBR Green (Applied Biosystems) and QuantiTect Primer Assays (QIAGEN).

**Statistics**

For data from the biobank binary logistic regression was used to examine the relationships between pulmonary fibrosis and chronotype, shift work or sleep duration. Both unadjusted and adjusted analyses were performed taking into account the known epidemiological associations for pulmonary fibrosis (age, sex and smoking)(15) as well as BMI.

Other data was evaluated using Student's t-test or two-way ANOVA for multiple comparisons as indicated. Significant values are shown as $p < 0.05$ (*), $p < 0.01$ (**).

**Animals**

All animals were maintained in 12h:12h light:dark (LD) with food and water supplied ad libitum. mPER2::luc transgenic mice were previously described (16). The Rev-erbα$^{fl/fl}$ mouse (Rev-erbαDBD$^{m}$) and Cre drivers targeting club cells (CCSP$^{icre}$) and myeloid cells (Lysm$^{cre}$) are as previously described (14). The PDGFRβ$^{cre}$ mouse was a kind gift from Henderson and has been previously described (17). The Bmal1$^{fl/fl}$ mouse has been previously described (9).

**Cell Lines**

mLF-hT cells have been recently described (18) and were a kind gift from Hinz. Briefly the primary murine lung fibroblasts had telomerase reverse transcriptase virally introduced preventing them from becoming senescent. Human lung fibroblast Mrc5 cells were purchased from the ATCC.

**Chemicals**

Bleomycin (sulfate from *Streptomyces verticillus*) was obtained from Sigma-Aldrich. The REVERB agonist, GSK 4112 (19) has been previously described and were obtained from Sigma.

**Study Approval**

Ethical approvals for both human and animal studies were obtained. The UK Biobank study was approved by the National Health Service National Research Ethics Service (ref. 11/NW/0382), use of human tissue was approved following permission from the Health Research Authority (11/NE/0291) and (14/NW/0260). All participants gave informed consent.

Animal procedures were carried out in accordance with the Animals Scientific Procedures Act (1986).

### Supplementary References


1. Allen N, *et al.* (2012) UK Biobank: Current status and what it means for epidemiology. *Health Policy and Technology* 1(3):123-126.
2. Herbert A, Wijlaars L, Zylbersztejn A, Cromwell D, & Hardelid P (2017) Data Resource Profile: Hospital Episode Statistics Admitted Patient Care (HES APC). *International journal of epidemiology* 46(4):1093-1093i.
3. Lane JM, *et al.* (2016) Genome-wide association analysis identifies novel loci for chronotype in 100,420 individuals from the UK Biobank. *Nature communications* 7:10889.
4. Mongrain V, Lavoie S, Selmaoui B, Paquet J, & Dumont M (2004) Phase relationships between sleep-wake cycle and underlying circadian rhythms in Morningness-Eveningness. *Journal of biological rhythms* 19(3):248-257.
5. Lane JM, *et al.* (2017) Genome-wide association analyses of sleep disturbance traits identify new loci and highlight shared genetics with neuropsychiatric and metabolic traits. *Nature genetics* 49(2):274-281.
6. Barrett T, *et al.* (2013) NCBI GEO: archive for functional genomics data sets--update. *Nucleic acids research* 41(Database issue):D991-995.
7. Yu G, *et al.* (2018) Thyroid hormone inhibits lung fibrosis in mice by improving epithelial mitochondrial function. *Nature medicine* 24(1):39-49.
8. Wynn TA, *et al.* (2011) Quantitative assessment of macrophage functions in repair and fibrosis. *Current protocols in immunology* Chapter 14:Unit14 22.
9. Gibbs J, *et al.* (2014) An epithelial circadian clock controls pulmonary inflammation and glucocorticoid action. *Nat.Med.* 20(8):919-926.
10. Liu H, Shah S, & Jiang W (2004) On-line outlier detection and data cleaning. *Computers & Chemical Engineering* 28(9):1635-1647.
11. Yamazaki S & Takahashi JS (2005) Real-time luminescence reporting of circadian gene expression in mammals. *Methods in enzymology* 393:288-301.
12. Smith-Clerc J & Hinz B (2010) Immunofluorescence detection of the cytoskeleton and extracellular matrix in tissue and cultured cells. *Methods in molecular biology (Clifton, N.J.)* 611:43-57.
13. Schindelin J, *et al.* (2012) Fiji: an open-source platform for biological-image analysis. *Nature methods* 9(7):676-682.
14. Pariollaud M, *et al.* (2018) Circadian clock component REV-ERBalpha controls homeostatic regulation of pulmonary inflammation. *The Journal of clinical investigation* 128(6):2281-2296.
15. Olson AL, Gifford AH, Inase N, Fernandez Perez ER, & Suda T (2018) The epidemiology of idiopathic pulmonary fibrosis and interstitial lung diseases at risk of a progressive-fibrosing phenotype. *European respiratory review : an official journal of the European Respiratory Society* 27(150).
16. Yoo SH, *et al.* (2004) PERIOD2::LUCIFERASE real-time reporting of circadian dynamics reveals persistent circadian oscillations in mouse peripheral tissues. *Proceedings of the National Academy of Sciences of the United States of America* 101(15):5339-5346.
17. Henderson NC, *et al.* (2013) Targeting of alphav integrin identifies a core molecular pathway that regulates fibrosis in several organs. *Nature medicine* 19(12):1617-1624.

18. Lodyga M*, et al.* (2019) Cadherin-11-mediated adhesion of macrophages to myofibroblasts establishes a profibrotic niche of active TGF-beta. *Science signaling* 12(564).
19. Meng QJ*, et al.* (2008) Ligand modulation of REV-ERBalpha function resets the peripheral circadian clock in a phasic manner. *J Cell Sci.* 121(Pt 21):3629-3635.
